# Histone Deacetylase 7 Mediates Tissue-Specific Autoimmunity via Control of Innate Effector Function in Invariant Natural Killer T-Cells

**DOI:** 10.1101/205377

**Authors:** Herbert G. Kasler, Intelly S. Lee, Hyung W. Lim, Eric Verdin

**Affiliations:** Gladstone Institute of Virology and Immunology, San Francisco CA 94158, USA; Department of Medicine, University of California, San Francisco, San Francisco CA 94158, USA; Buck Institute for Research on Aging, Novato, CA 94945, USA

## Abstract

We report that Histone Deacetylase 7 (HDAC7) controls the thymic effector programming of Natural Killer T (NKT) cells, and that interference with this function contributes to tissue-specific autoimmunity. Gain of HDAC7 function in thymocytes blocks both negative selection and NKT development, diverting these cells into a Tconv-like lineage. Conversely, HDAC7 deletion promotes thymocyte apoptosis and causes aberrant expansion of innate-effector cells. Investigating the mechanisms involved, we found that HDAC7 binds PLZF and modulates PLZF-dependent transcription. Moreover, HDAC7 and many of its transcriptional targets are human risk loci for IBD and PSC, autoimmune diseases that strikingly resemble the disease we observe in HDAC7 gain-of-function in mice. Importantly, reconstitution of iNKT cells in these mice abrogated their disease, suggesting that interaction between the defects in negative selection and iNKT cells caused by altered HDAC7 function can cause tissue-restricted autoimmunity, a finding that may explain the association between HDAC7 and hepatobiliary autoimmunity.

## Introduction

To become mature T cells, thymocytes must navigate through a complex process of selection and instruction, centered around signals received through their newly created T cell antigen receptors (TCRs). For thymocytes destined to become conventional naïve CD4 or CD8 T cells (Tconv), this requires passing two key checkpoints: positive selection, in which cortical CD4/CD8 double-positive (DP) thymocytes must receive a minimum level of TCR stimulation from self peptide-MHC complexes in order to adopt the appropriate lineage and continue maturation, and negative selection, in which thymocytes with self-reactivity above a critical threshold are deleted from the repertoire by activation-induced apoptosis. While the elucidation of these mechanisms decades ago established a basic conceptual framework for the creation of a competent and self-tolerant T cell repertoire, the years since have brought to light an ever-increasing variety of alternate developmental programs that produce specialized populations of mature T cells functionally distinct from Tconv. These populations, critical for both effective host defense and self-tolerance, are elicited from the diverse pool of T cell precursors by specialized selection mechanisms, mostly involving strong recognition of noncanonical ligands, as in the case of NKT cells (Kronenberg 2014), or recognition of peptide-MHC ligands at high TCR avidities near the threshold of negative selection, as in the case of nTreg or CD8αα IEL (Klein et al. 2014, Moran et al. 2011). The TCR signals involved in their development are generally stronger than those that mediate positive selection to the Tconv lineage, and the process is thus termed agonist selection (Stritesky, Jameson, and Hogquist 2012).

One feature that distinguishes many of these specialized cell types from Tconv is the thymic acquisition of constitutive effector function, a phenotype shared with innate immune cells and thus giving rise to the term “innate-like” or “innate effector” T cells. Whereas Tconv exit the thymus with a naïve phenotype, circulate broadly, and require a several days long, orchestrated process of priming and clonal expansion to become fully functional effector/memory cells, innate-like T cells are often constitutively tissue-resident and make mature effector responses to their cognate stimuli immediately (Kang and Malhotra 2015). Innate-like T cells exit the thymus larger than Tconv, with an antigen-experienced phenotype and an expanded secretory apparatus, allowing them to rapidly elaborate robust cytokine responses after brief TCR stimulation (Kang and Malhotra 2015, Chandra and Kronenberg 2015, Brennan, Brigl, and Brenner 2013). These differences arise due an alternative thymic maturation process that parallels the priming of naïve T-cells in the periphery. For NKT cells, this includes a ~100-fold intra-thymic proliferative expansion to generate pre-established clonal populations (Benlagha et al. 2002). Maintenance of the innate effector phenotype in NKT cells can at least partially be attributed to stable expression of their signature transcription factor Promyelocytic Leukemia Zinc Finger Protein (PLZF, ZBTB16) (Kovalovsky et al. 2008, Savage et al. 2008). PLZF expression is established during thymic development of NKT cells, via a cellular mechanism that involves strong recognition of glycolipid ligands on the non-canonical CD1D MHC molecule by a clonally restricted (in mice, Vα14/Jα18 with one of several possible β chains) TCR, together with homotypic co-stimulation through the SAP family of co-receptors (Bendelac, Savage, and Teyton 2007). However, what downstream factors link these surface signals to stable PLZF expression and what other pathways may be involved are still open questions.

We have previously described how Tconv development is regulated by the class IIA histone deacetylase Histone Deacetylase 7 (HDAC7), a TCR signal-regulated corepressor abundantly expressed in thymocytes (Dequiedt et al. 2003, Kasler and Verdin 2007). The activity of HDAC7 is controlled by nuclear exclusion in response to phosphorylation of conserved serine residues in their N-terminal adapter domains (Verdin, Dequiedt, and Kasler 2003). In thymocytes, TCR stimulation results in HDAC7 phosphorylation and nuclear exclusion via protein kinase D (Parra et al. 2005). CD4/CD8 double-positive (DP) thymocytes lacking HDAC7 are much more likely than WT thymocytes to die before becoming positively selected, significantly impeding their development into mature Tconv (Kasler et al. 2011). Conversely, if a transgene encoding a phosphorylation-deficient, constitutively nuclear version of HDAC7 (HDAC7ΔP) is transiently expressed in the thymus at sub-endogenous levels (Kasler et al. 2012), deletion of autoreactive thymocytes by negative selection is strongly blocked and the hosts develop lethal autoimmunity. Consistent with broad blockade of negative selection, we observed autoantibodies to a comprehensive array of tissue antigens in these mice. However, for reasons that were not clear to us at the time actual tissue destruction occurred almost exclusively in a gastrointestinal/hepatobiliary compartment that is anatomically tied together by the contiguous epithelial surfaces of the GI lumen and the pancreatic and biliary ductal systems (Kasler et al. 2012).

The potential significance of this peculiar pattern of HDAC7-mediated autoimmunity for human disease has recently been brought into sharp focus by two separate studies identifying polymorphisms at the loci of HDAC7 as well as several of its upstream regulatory kinases as independent risk factors in human inflammatory bowel disease (IBD), and also in primary sclerosing cholangitis (PSC), a destructive autoimmune syndrome of the hepatobiliary system, which is additionally associated with increased IBD risk (Liu et al. 2013, Jostins et al. 2012). The striking parallel between these human syndromes and the autoimmunity observed in HDAC7ΔP transgenic mice suggested to us a connection between HDAC7 and these types of autoimmunity that goes beyond simply blocking thymic negative selection. This led us to undertake a more thorough phenotypic characterization of mice with altered HDAC7 function during T cell development, revealing that HDAC7 has a key role in the regulation of the innate effector programming of iNKT cells, at least in part via direct modulation of the transcriptional activity of PLZF. Both gain and loss of HDAC7 function in thymocytes resulted in aberrant effector programming of T cells in both the Tconv and innate-like lineages, leading to multiple abnormalities in peripheral populations. These studies shed new light on the molecular pathways that regulate the effector programming of innate-like T cells, reveal a new key molecular target of HDAC7 in T cell development, and set forth a novel cellular model of autoimmunity in which one genetic lesion mediates multiple defects in thymic selection, which converge in the periphery to produce a unique, tissue-restricted pattern of disease. Given the established genetic association between HDAC7 and very similar human syndromes, our findings are likely to be of considerable significance in the understanding of these diseases.

## Results

### Alteration of HDAC7 Function Dysregulates Thymic Innate Effector Programming and Interferes With iNKT Development

We previously showed that if a constitutively nuclear mutant of HDAC7 (HDAC7ΔP) is transiently expressed at normal levels during thymic T cell development but not in mature T cells, autoreactive cells that would normally die by negative selection instead exit the thymus as naïve Tconv (Kasler et al. 2012). However in our previous study we did not assess the fates of cells destined to become innate effectors. Analyzing these populations, we noted a modest suppression of Treg and CD8αα IEL (not shown), but the most striking observation we made was the near total absence of invariant Natural Killer T cells (iNKT), an oligoclonal population that is reactive to α-galactosylceramide (αGalCer) presented by the CD1D non-canonical MHC molecule (CD1D/αGalCer) (Kronenberg 2014). Cells positive for staining with CD1D/αGalCer tetramers represent approximately 3% of TCRβ-positive cells in wild type C57BL/6 (B6) thymus and 30% in liver, however they are nearly undetectable in either of these tissues or in the spleens of HDAC7ΔP mice (Fig. 1A, B, Fig. S1A, B), suggesting a profound, early block in iNKT development. The few tetramer-binding cells remaining in HDAC7ΔP mice were CD24lo, but failed to upregulate either CD44 or NK1.1 (Fig. 1C), indicating a Stage 1 block according to the conventional staging system for iNKT development (Stritesky, Jameson, and Hogquist 2012). We also evaluated the prevalence of CD44/NK1.1-expressing T cells that were not tetramer-reactive, and noted a marked reduction in their numbers as well (Fig. S1E-F), indicating a broad defect in the development of the NKT lineage.

**Figure 1.**
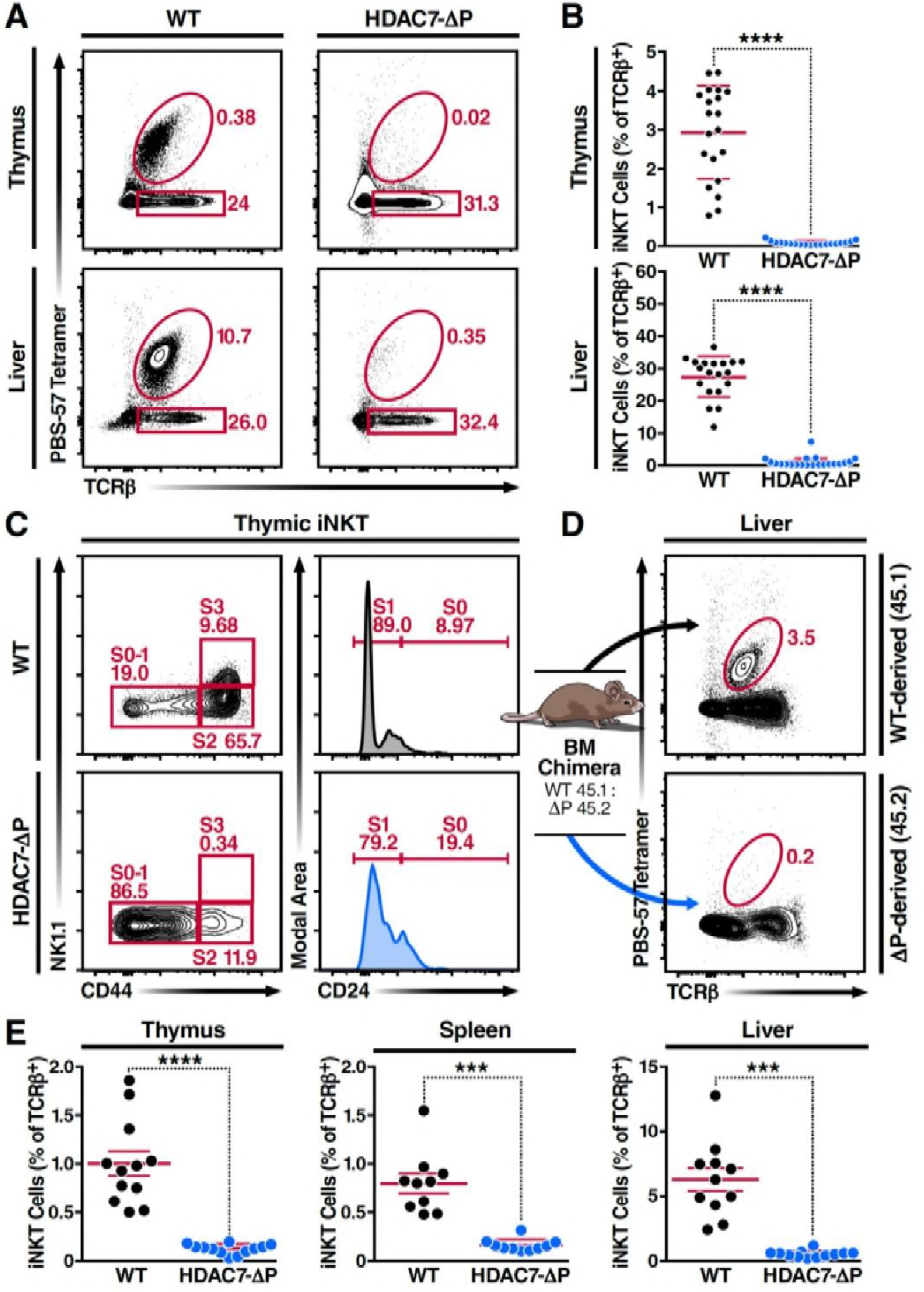
A Gain-of-Function HDAC7 Mutant, HDAC7ΔP, Arrests Thymic iNKT Development. (A, B) Flow cytometric plots (A) and quantification (B) of iNKT cells and conventional αβ T-cells identified by staining with TCRβ and PBS-57-loaded tetramer (“Tet”) in thymus and liver. iNKT cells are Tet^+^ TCRβ^+^, while conventional T-cells are Tet^−^ TCRβ^+^. (C) Conventional staging of iNKT development based on CD24, CD44 and NK1.1 expression in Tet^+^ TCRp^+^ thymic iNKT cells. (D, E) Representative flow cytometric plots (D) and total quantification (E) of iNKT cells in irradiated mixed WT (CD45.1): *HDAC7ΔP* (CD45.2) bone-marrow chimeras from thymus, spleen and liver. Bars on graphs indicate mean ± SEM (error bars); symbols represent individual mice. Data in (B) are combined from 8 independent experiments involving 1-3 littermate pairs; data in (C) are representative of 3 experiments with 3-5 mice per group. Data in (D-E) are representative of 3 experiments with 4-6 mice per group. Statistical significance was determined using unpaired two-tailed t tests; ***p ≤ 0.001, ****p ≤ 0.0001.

To rule out cell-extrinsic mechanisms for this block, we generated mixed hematopoietic chimeras reconstituted with a 1:1 mixture of wild-type (WT) and HDAC7ΔP bone marrow. As we previously reported (Kasler et al. 2012), the HDAC7ΔP transgenic population contributed robustly to the pool of CD4 and CD8 SP thymocytes, although there was a transient reduction in prevalence at the immature single positive (ISP) stage (Fig. S1C). At early time points post-reconstitution (6-8wk), the distributions of naïve and memory T-cells in peripheral CD4+ and CD8+ Tconv subsets were equivalent as well (Fig S1D). However, while the wild type-derived population reconstituted hepatic iNKT cells efficiently, HDAC7ΔP bone marrow failed to contribute significantly to this compartment (Fig. 1D, E). This was also true in the thymus and spleen (Fig. 1E), demonstrating that the block observed in the intact transgenic mice was due to a cell-autonomous mechanism.

We next examined the effects of loss of HDAC7 in the thymus on these phenotypes, using our previously characterized strain that deletes loxp-flanked HDAC7 under the control of the p56lck proximal promoter (HDAC7 flox/lck-cre, henceforth HDAC7 KO) (Kasler et al. 2011). We previously reported that loss of HDAC7 during T cell development increased apoptosis of DP thymocytes leading to inefficient positive selection. This shortened thymocyte lifespan resulted in a truncation of the TCR Jα repertoire, with distal rearrangements underrepresented (Kasler et al. 2011). It was thus not surprising to find that HDAC7 KO mice with an endogenous TCR repertoire had fewer iNKT cells than WT controls; for example, HDAC7 KO liver contained 80-90% fewer iNKT cells as a proportion of TCRβ+ T-cells (Fig. 3A). This roughly five-fold reduction, consistent with the degree of underrepresentation of the relatively distal Jα18 TCR segment we previously noted (Kasler et al. 2011), was similarly observed in the spleen and thymus (Fig. S3B). Importantly, unlike the residual tetramer-reactive cells in HDAC7ΔP mice, iNKT calls in HDAC7 KO mice had normal expression of CD44 and NK1.1, suggesting that their development was not functionally altered. (Fig, 2B)

**Figure 2.**
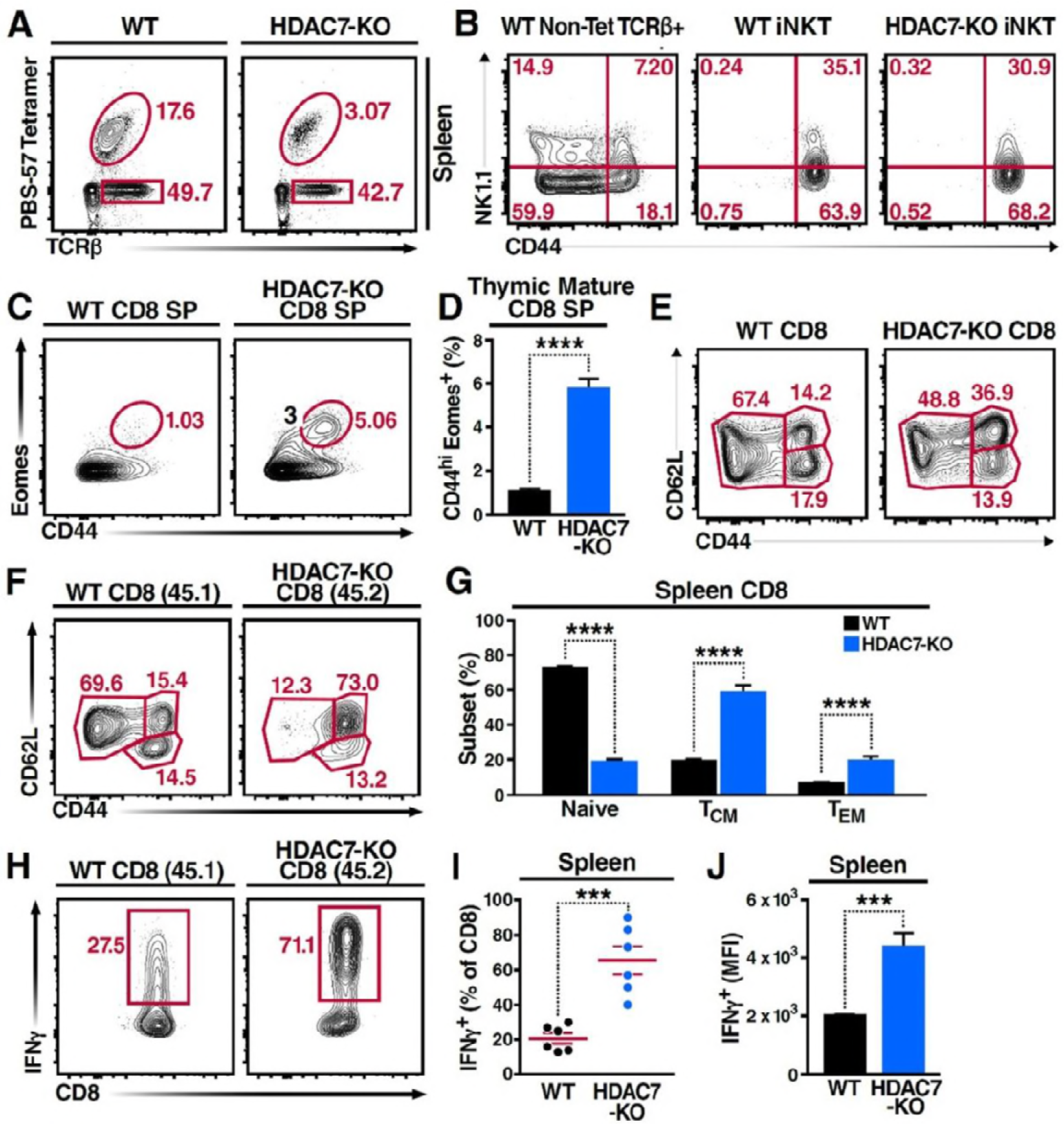
Deletion of HDAC7 in thymocytes Reduces iNKT Numbers and Expands an Innate-Memory CD8 Population. (B) Representative flow cytometric staining of CD44 and NK1.1 from peripheral iNKT (Tet^+^ TCRβ^+^) T-cells compared with non-tetramer binding (“Non-tet”) TCRβ^+^ T-cells. Note that HDAC7-KO iNKT cells are phenotypically normal. (C, D) Identification of an expanded CD44^hi^ Eomes^+^ innate memory population in CD8 thymocytes from HDAC7 KO mice. Representative plots shown in (C) with quantification shown in (D) for littermates. Mature CD8 SP thymocytes are identified as TCRβ^+^CD8^+^CD4^−^. (E) Expression of CD44 and CD62L in CD8 T-cells from spleens of separate WT and HDAC7-KO age-and sex-matched littermate mice. Data are representative of 3 independent experiments with N = 2-4 mice per group. Representative flow plots (F) and total quantification (G) of peripheral naive, central memory (T_CM_), and effector memory (T_EM_) CD8 T-cell populations from WT (CD45.1) and *HDAC7 KO* (CD45.2) derived bone marrow in irradiated mixed BM chimeras. (G, H). Representative flow plots (H) and total quantification (I) of IFNγ secretion in *ex vivo* stimulated CD8 T-cells. Splenocytes were harvested from mixed WT (CD45.1) / *HDAC7 KO* (CD45.2) BM chimeras, stimulated *ex vivo* for 4h with PMA/Ionomycin. (J) Median fluorescence intensity (MFI) of IFNγ+ secretion in ex vivo-stimulated CD8 T-cells from (I). Bars on graphs indicate mean ± SEM (error bars); symbols represent individual mice. Data in (D, I, J) are combined from 3 independent experiments with at least 3 mice per group; data in (G) are combined from 4 independent experiments with at least 2 mice per group. Statistical significance was determined using either unpaired two-tailed T-test (D, I, J) or two-way ANOVA (G); ***p ≤ 0.001, ****p ≤ 0.0001. A Bonferroni post-test was used for pairwise comparisons in (G).

**Figure 3.**
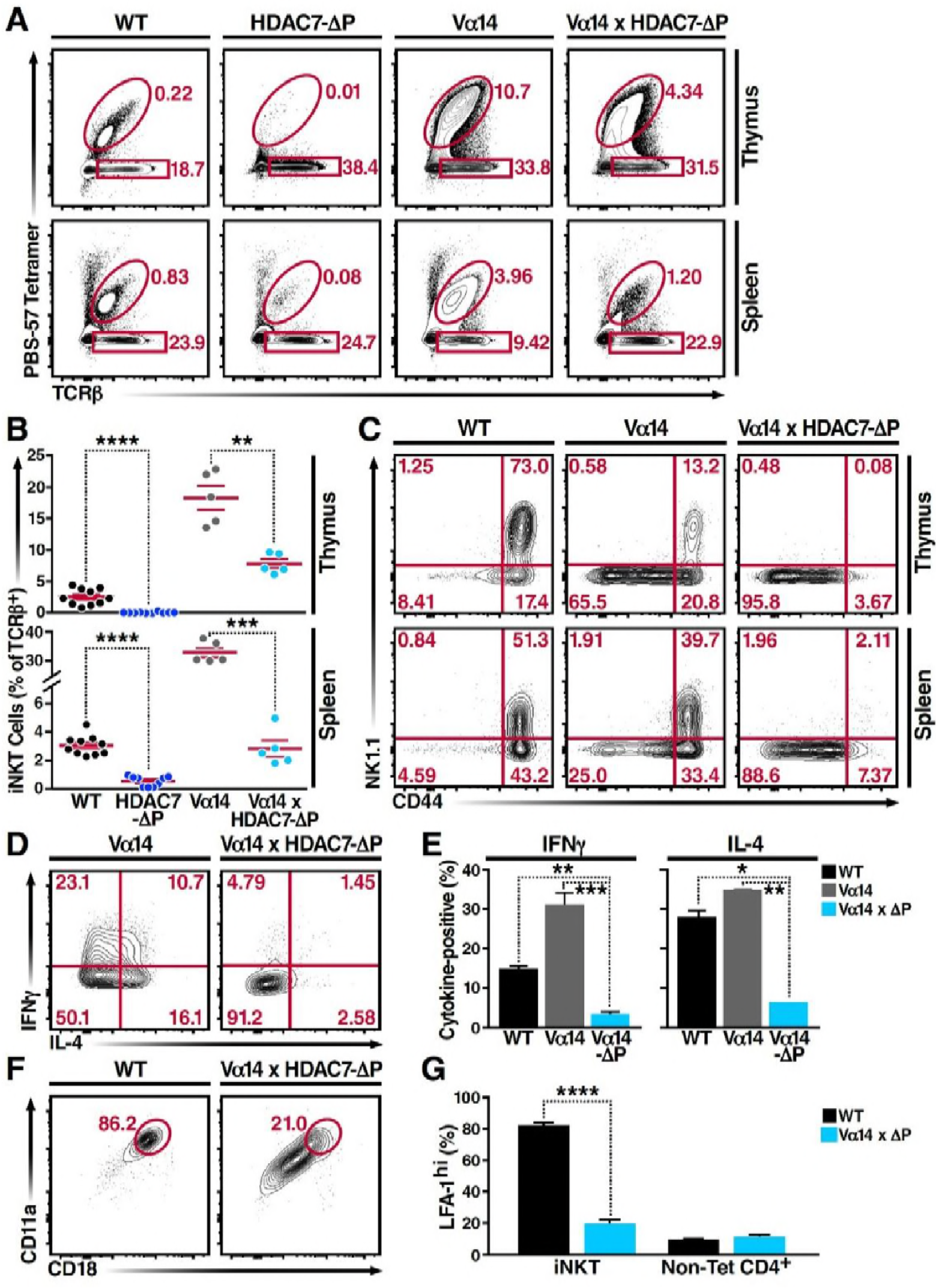
HDAC7ΔP Blocks Innate Effector Development in iNKT Cells and Converts Them to Naive-Like T-cells. (A, B) Flow cytometric plots (A) and total quantification (B) of iNKT cells (Tet^+^ TCRβ^+^) from thymus (top) and spleen (bottom) of littermate mice with the indicated genotypes. (C) Representative surface expression of CD44 and NK1.1 in iNKT cells from thymus and spleen in littermate mice of the indicated genotypes, based on mice from (B). (D, E) Representative staining (D) and total quantification (E) of IFNγ and IL-4 secretion in total splenocytes from littermate mice of the indicated genotypes, stimulated *ex vivo* for 4h with PMA/Ionomycin. (F) Surface expression of LFA-1 (CD11a/CD18) in splenic iNKT (Tet^+^/TCRβ^+^) cells. Bars on graphs indicate mean ± SEM; plot symbols represent individual mice. Data in (B) are from 4 independent experiments; data in (E, G) are from 3 independent experiments. Statistical significance was determined using one-way (B, E) or two-way (G) ANOVA; *p ≤ 0.05, **p ≤ 0.01, ***p ≤ 0.001, ****p ≤ 0.0001. Tukey (B, E) or Bonferroni post-tests (G) were used for pairwise comparisons.

Although deletion of HDAC7 did not result in expansion of NK1.1-expressing T cells, we did observe significant abnormalities in the effector programming of non-tetramer-reactive thymocytes. We noted a substantial expansion of a CD44hi Eomes+ population in the mature CD8 SP compartment in the thymus (Fig. 2C-D). Examination of the peripheral CD8 T cells in these animals also showed a substantial increase in CD44 expression, suggesting an expansion of innate effector CD8 cells (Fig. 2E). These cells resemble Eomes+ innate memory CD8+ cells that are typically generated *in trans*, in response to IL4 secretion by thymic-resident iNKT cells (Lee et al. 2013, Weinreich et al. 2010), however as previously noted iNKTcells are depleted rather than expanded in HDAC7 KO mice, suggesting a different mechanism. To clarify this question, we examined the phenotypes resulting from loss of HDAC7 in WT: HDAC7 KO mixed hematopoietic chimeras.

In 1:1 chimeras, HDAC7 KO thymocytes competed equally through the ISP stage, but thereafter competed poorly and became steadily less abundant. This substantial underrepresentation of HDAC7 KO CD4SP and mature CD8SP thymocytes in 1:1 chimeras (Fig. S2A-B) made analysis at this stage difficult, however their representation in the periphery was sufficient. In the spleen, we saw a strong increase in CD44 expression in the HDAC7 KO-derived vs. to the WT-derived CD8 T cell population (Fig. 2F-G), duplicating what we saw in the intact mice and indicating that the phenotypes we observed are likely cell-autonomous. To further characterize the phenotype of these cells, we briefly stimulated splenocytes from these chimeras *ex vivo*, and found that HDAC7 KO-derived CD8+ T-cells produced much more IFNγ than WT-derived CD8+ T-cells in the same culture, assessed both as percent cytokine-positive (Fig. 3H, I) and by median fluorescence intensity (MFI) of cytokine staining (Fig. 3J). CD8+ T-cells from HDAC7 KO population also had increased expression of the Eomes-associated chemokine receptor CXCR3 and the trafficking receptor Ly6C (Fig. S2C).

Loss of HDAC7 thus appears to result in the aberrant adoption of NKT-like innate effector programming by CD8 SP thymocytes that would otherwise have exited the thymus as naive Tconv. We observed a much more modest degree of abnormality in the CD4 compartment (not shown), which we hypothesize is due to the greater similarity that CD8 thymic selection bears to NKT selection, in terms of both the similarity of CD1D to Class I MHC and the availability of selecting ligands on all thymocytes rather than just on specialized thymic APC. Loss of HDAC7 may thus allow some DP thymocytes to aberrantly adopt this lineage through some partial analogue of NKT selection. Collectively, our findings with both the HDAC7 knockout and HDAC7ΔP transgene findings suggest that HDAC7 may function as a gatekeeper of innate effector programming, blocking the functional maturation of iNKT cells when constitutively expressed in the nucleus, and conversely allowing the aberrant acquisition of innate effector characteristics in CD8 T cells when it is conditionally deleted.

### HDAC7 Regulates the Effector Programming of NKT Cells in a Manner That Mirrors the Function of PLZF

To generate a larger population of iNKT precursors for more in-depth evaluation the role of HDAC7, we employed the Vα14-Jα18 TCRα transgene (henceforth “Vα14”), encoding the invariant TCR α chain that when paired with the appropriate endogenous β chains allows iNKT cells to bind glycolipids with high affinity (Griewank et al. 2007). Expressing this TCR transgene greatly increases the frequency of CD1D/αGalCer-reactive thymocytes, which arise naturally only at only around 1 in 10^4^ cells. As expected, mice expressing only the Vα14 transgene had many more iNKT cells in thymus and spleen than WT mice (Figs. 3A-B and S3A-B). Also consistent with our expectations, when we crossed the Vα14 TCRα transgene into the HDAC7 KO strain, we observed a complete rescue of iNKT cell abundance in the thymus and periphery (Fig. S3A-B), resulting in identical numbers between Vα14 and Vα14 HDAC7 KO mice. These cells were phenotypically similar to WT iNKT cells in terms of CD44/NK1.1 expression and cytokine production (not shown), suggesting that shortened thymocyte lifespan was indeed the key cause of the lowered iNKT abundance in HDAC7 KO mice.

In contrast to this finding, when the Vα14 transgene was co-expressed with HDAC7ΔP, the rescue in the numbers of CD1D/αGalCer-reactive cells was incomplete (Fig. 3A-B), and the cells were phenotypically abnormal (Fig. 3, S3). This result suggests that rather than blocking the maturation of CD1D/αGalCer-reactive cells categorically, HDAC7ΔP blocked the intrathymic proliferation that is normally associated with post-positive selection iNKT differentiation (Benlagha et al. 2002), directing the cells instead to mature without proliferating, as if they were positively selected Tconv. Consistent with this idea, other characteristics of CD1D/αGalCer-reactive Vα14 × HDAC7ΔP T cells were similar to those of naïve Tconv. Flow analysis revealed that like the residual tetramer-reactive cells present in the HDAC7ΔP mice (Fig. 1C) the rescued iNKT cells in Vα14 × HDAC7ΔP mice failed to upregulate the memory marker CD44 or the NKT marker NK1.1 in the thymus like their Vα14-only counterparts (Fig. 3C, top row). This phenotype persisted in the spleen, after the HDAC7ΔP transgene was turned off (Fig. 2C, bottom row), suggesting that the cells had failed to undergo effector programming in the thymus.

We next examined their cytokine responses to brief *ex-vivo* stimulation. When stimulated for 4 hours with PMA/ionomycin, CD1D/αGalCer-reactive WT and Vα14 transgenic iNKT cells exhibited a robust cytokine response, secreting both IFNγ and IL-4. In contrast, Vα14 × HDAC7ΔP iNKT were far less likely to make IFNγ or IL-4 (Fig. 3D-E), as would be expected for naïve Tconv. Additionally, iNKT cells typically express high levels of the integrin LFA-1 (CD11a/CD18), allowing them to remain localized in tissue-specific vascular beds such as hepatic sinusoids (Thomas et al. 2011). In contrast, Vα14 × HDAC7ΔP iNKT cells exhibited far lower expression levels (Fig. 3F-G), comparable to those seen in circulating non-CD1D/αGalCer-reactive CD4+ (mainly naïve) T-cells (Fig. 3G, right). Moreover, while Vaα14 × HDAC7ΔP iNKT cells were found at comparable frequency in spleen to WT iNKT cells, they failed to concentrate in peripheral tissues such as the liver (Fig. S3C-D), a behavior more characteristic of naïve Tconv rather than iNKT cells. Thus, constitutively nuclear HDAC7, encoded by HDAC7ΔP, prevents iNKT precursors from initiating innate effector development. As they have low CD44 expression, produce few cytokines after brief stimulation, and freely recirculate, they appear to become diverted into functionally naïve-like T-cells.

When considering how both gain and loss of thymic HDAC7 function alter innate effector development, we were struck by how closely our results mirrored findings reported in similar studies of the transcription factor PLZF. Notably, the prominent Stage 1 iNKT block and loss of effector memory phenotype in residual peripheral iNKT cells observed in gain-of-function HDAC7ΔP strongly resembles the iNKT defect observed in PLZF knockouts (Kovalovsky et al. 2008, Savage et al. 2008). Conversely, the consequences of loss of HDAC7 function – notably expansion of Eomes+ innate memory CD8+ coupled with the appearance of IFNγ-secreting memory CD8+ T-cells (Fig. 2C, H) mirror results reported in gain-of-function PLZF transgenic mice (Kovalovsky et al. 2010, Savage, Constantinides, and Bendelac 2011). Polyclonal (noninvariant) type II NKT cells are also thought to be PLZF-dependent (Zhao et al. 2014), and we similarly noted a near absence of tissue-resident type II NKTs in HDAC7ΔP mice, defined by a Tet-TCRβ+CD8-CD44hiNK1.1+ profile (Fig. S1E-F). HDAC7 and PLZF thus appear to play nearly inverse roles in iNKT development (Fig. 4E).

**Figure 4.**
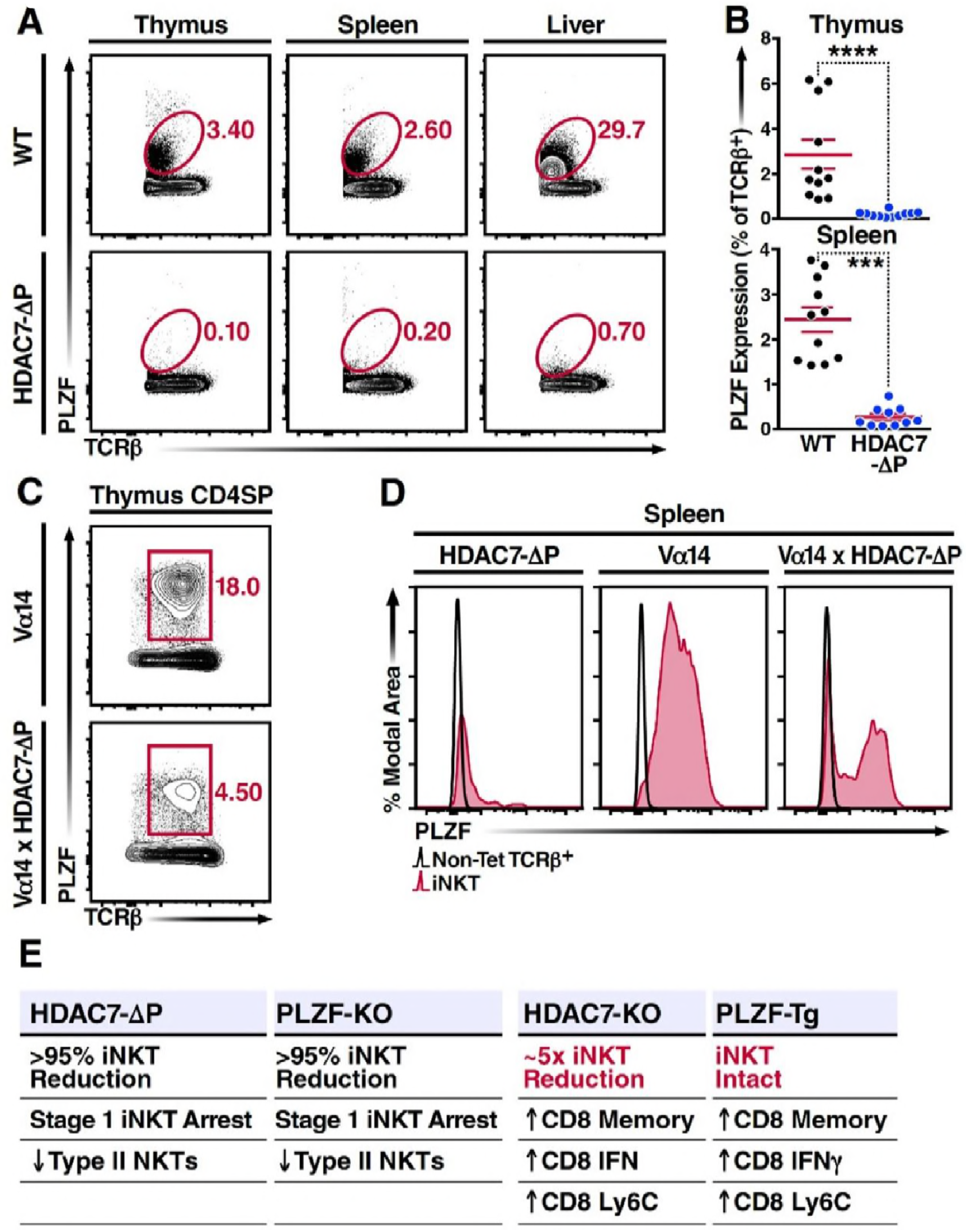
Nuclear HDAC7 Retention Restricts PLZF Expression and Mirrors PLZF-Associated T Cell Phenotypes. (A, B) Representative flow cytometric plots (A) and total quantification (B) of PLZF expression in TCRβ^+^ cells from thymus, spleen, and liver. (C) PLZF expression in mature CD4 SP (CD4^+^ CD8^−^ TCRβ^+^) thymocytes. (D) PLZF expression in peripheral iNKT (Tet^+^ TCRβ^+^) cells from spleen. Black unfilled histograms correspond to conventional (Tet^−^TCRβ^+^) T-cells, red tinted to iNKT (Tet^+^TCRβ^+^) cells. (E) Summary table comparing phenotypes in HDAC7ΔP, PLZF KO, HDAC7 KO and PLZF Tg mice with respect to iNKT and conventional T-cell development. Bars on graphs in (B) indicate mean ± SEM; symbols represent individual mice. Data in (B) are combined from 4 independent experiments with at least 2 mice per group; data in (D) are representative of 3 independent experiments with 2 mice per group; Statistical significance was determined using unpaired two-tailed T-tests (B); ***p ≤ 0.001, ****p ≤ 0.0001 vs. WT.

One possible mechanism for this inverse relationship is the idea that nuclear HDAC7 represses the expression of PLZF, preventing HDAC7ΔP thymocytes from expressing PLZF (Seiler et al. 2012). Indeed, we observed a pronounced reduction in PLZF expression in TCRβ+ T-cells from HDAC7ΔP mice in all organs examined, including thymus, spleen and liver (Fig. 4A, B). However, robust PLZF was still detected in CD4+ SP thymocytes from Vα14 × HDAC7ΔP mice; although expression was restricted compared to Vα14-only thymus (Fig. 4C), Interestingly, PLZF expression was maintained in roughly half of splenic Vα14 × HDAC7ΔP iNKT cells (Fig. 4D, right panel). Thus, transcriptional repression of PLZF expression by HDAC7 is probably insufficient to fully explain the iNKT block, as even PLZF+ Vα14 × HDAC7ΔP iNKT cells remain naïve-like, Stage 1-arrested T-cells (Fig. 3C).

### HDAC7 and PLZF Inversely Regulate a Shared Innate Effector Gene Network That is Highly Relevant to Autoimmune Disease

The remarkable inverse relationship between the phenotypes mediated by alterations of HDAC7 and PLZF function in iNKT cell development prompted us to take an unbiased, genome-wide approach to understanding how these two factors might coordinately regulate the transcriptional landscape of this process. To this end, we generated gene expression profiles by RNA-seq of Cd1d/αGalCer-reactive Vα14 Tg and Vα14 X HDAC7ΔP Tg thymocytes and splenocytes, as well as polyclonal naïve conventional CD4 SP thymocytes and splenocytes. Differential gene expression profiles were constructed for Vα14 Tg vs. naïve Tconv, Vα14 X HDAC7ΔP Tg vs naïve Tconv, and Vα14 X HDAC7ΔP vs. Vα14 Tg, by comparing the normalized scalar expression values for 3 biological replicates of each condition, based on roughly 40 million mapped reads per sample (See Table S1, Experimental Procedures). When we plotted significant expression changes for Vα14 Tg vs. Tconv in ether thymus or spleen (Fig. 5A, left and right panels, horizontal axes) against the corresponding changes for Vα14 X HDAC7ΔP vs Tconv (vertical axes), it was evident that HDAC7ΔP makes the overall gene expression pattern of Cd1d/αGalCer-reactive cells more similar to that of Tconv, as shown by the clockwise shift of the plot trend line from the diagonal in both tissues (Fig. 5A, solid plot diagonal vs. dotted trend line). Reflecting this effect, iNKT development-associated gene expression changes (both up and down) that were suppressed by HDAC7ΔP (Fig. 5A, green plot points and numbers) greatly outnumbered those enhanced by HDAC7ΔP (Fig. 5A, red plot points and numbers) in both spleen and thymus. Strongly induced genes involved in iNKT cell development that were suppressed by HDAC7ΔP included ID2, PLZF, NK1.1, T-bet, Gata3, IL4, IFNG, and Hobit, a zinc-finger transcription factor recently shown to be essential for the acquisition of tissue-resident effector function (Mackay et al. 2016) (Fig. 5A, labeled points). This pattern of suppression was established in the thymus (Fig. 5A, left), but clearly persisted in the spleen (Fig. 5A, right), after expression of HDAC7ΔP was turned off. Blocking HDAC7 nuclear export in the thymus thus apparently programs a more naïve-like state of differentiation into CD1D/αGalCer-reactive cells that persists even after HDAC7-mediated repression is removed.

**Figure 5.**
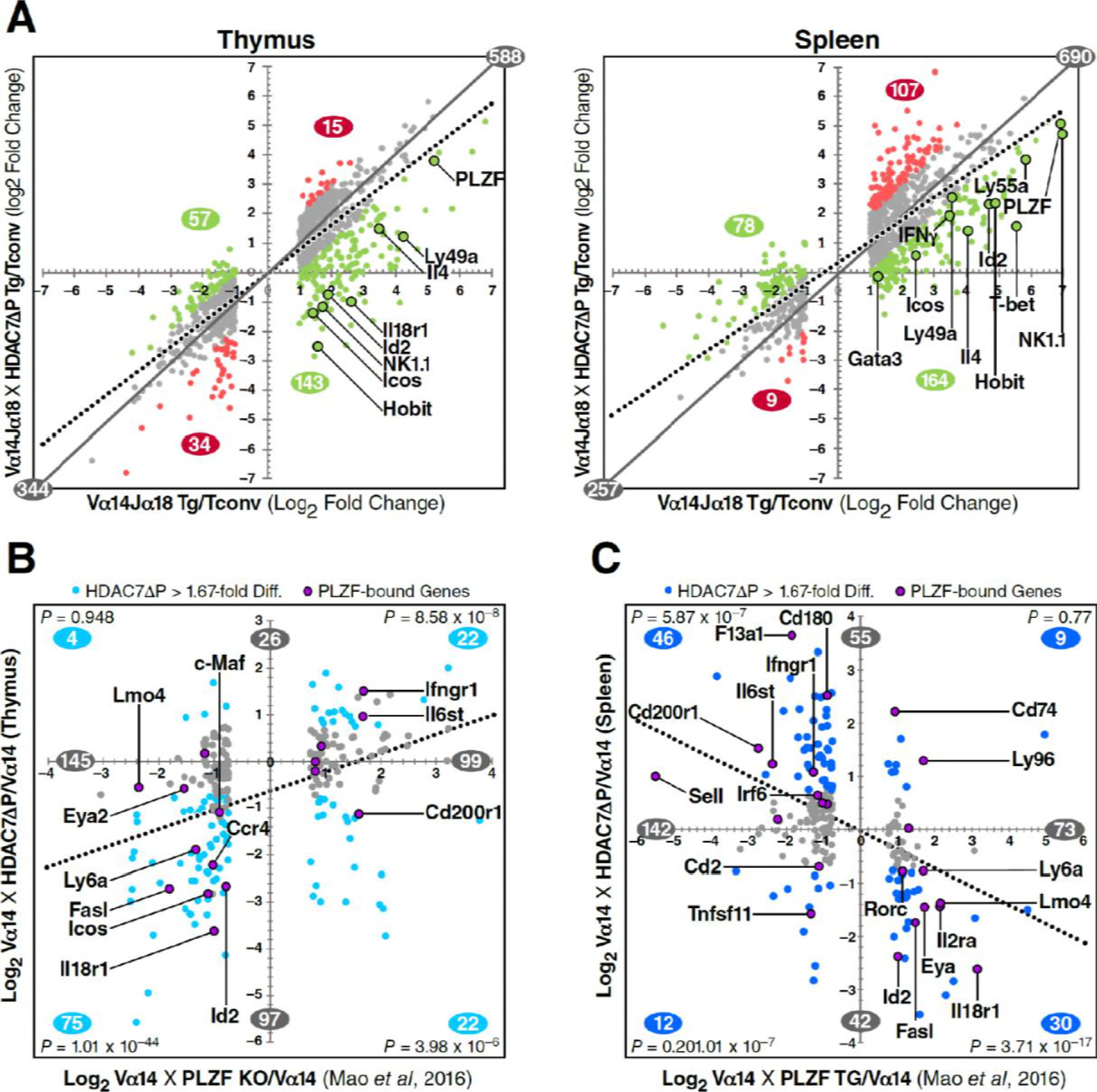
HDAC7 Regulates a Cassette of Genes in Glycolipid-Reactive Cells That is Highly Relevant to Innate Effector Function, Inflammation, Autoimmunity, and Autoimmune Liver Disease. (A) Scatter charts showing gene expression changes in Cdld/αGalCer-reactive *Vα14* Tg (X axis) and *Vα14* X *HDAC7ΔP* Tg (Y axis) thymocytes (left) or CD4 splenocytes (right) vs naïve CD4SP thymocytes or splenocytes, respectively. The solid gray line indicates the plot diagonal and the dotted gray line indicates the Least Squares best-fit line of the plotted data. Genes displayed were expressed at least 1.75-fold differentially between tetramer-reactive and naive cells, with p<0.05 (2-tailed Student’s T test) for 3 biological replicates of each genotype. Colored plot points represent genes whose differential expression vs. naive was enhanced (red points) or suppressed (green points) at least 1.75-fold by co-expression of *HDAC7ΔP* (C, D) Scatter charts showing genes >1.66-fold differentially expressed due to loss of PLZF function (C, horizontal axis), or due to transgenic expression of PLZF (D, horizontal axis) according to (Mao et al. 2016), plotted against effect of HDAC7AP expression in PBS-57 tetramer-reactive *Vα14* X *HDAC7ΔP* vs *Vα14* Transgenic thymocytes in Thymus (C, vertical axis) or spleen (D, vertical axis). Total number of genes >1.67-fold differentially expressed along each axis are indicated in gray. Numbers of genes, with P-values (binomial distribution) of the overlap, for genes differentially expressed along both axes in each quadrant (blue symbols), are indicated in blue.

These data were also helpful in identifying key candidate molecular targets of HDAC7. IPA analysis of putative upstream regulators of the HDAC7-affected gene set identified multiple targets highly relevant to iNKT development and function, including PLZF, ID2, IL4, IFNG, T-bet, and GATA3 (Fig. S4A, for a complete list of putative upstream regulators see Table S3). The downstream targets of these were almost universally affected in a manner that suggests inhibition rather than activation of the putative upstream regulator. The expression of most of these upstream regulators was itself suppressed by HDAC7, suggesting an obvious mechanism of regulation. However the Tec kinase ITK, the most highly correlated upstream regulator of HDAC7 targets in both thymus and spleen, was only modestly suppressed in spleen and not significantly suppressed in thymus, suggesting that HDAC7 might regulate its activation more than its expression. ITK has a well-characterized role in the maturation of conventional CD8 T cells, CD8 innate effectors, and iNKT cells (Atherly et al. 2006, Felices and Berg 2008).

Similarly, PLZF expression was only modestly repressed by HDAC7ΔP (Fig. 5A), yet its downstream targets were very highly correlated with the HDAC7 target gene set, based on both IPA analysis and comparison of HDAC7-regulated genes with genes identified in a recent, comprehensive study of PLZF-regulated genes in iNKT cell development (Mao et al. 2016) (Fig. 5B-C, Fig. S4C). Gene expression changes due to loss of PLZF function in iNKT cells show a clear positive correlation with changes caused by expression of HDAC7ΔP (Fig. 5B, Fig. S4C), while changes caused by expression of a PLZF transgene show a clear negative correlation (Fig. 5D, Fig. S4C), demonstrating an inverse relationship between HDAC7 and PLZF function. Genes that were found to associate directly with PLZF by chIP-seq (Mao et al. 2016), cluster strongly around the HDAC7-PLZF diagonals, and are highly concentrated among the most negatively correlated genes in terms of the effects of HDAC7 vs. PLZF function (Fig. 5D-E, labeled genes, Fig. S4C, red asterisks). Out of the 31 genes reported by Mao, et al. to be both bound by PLZF and differentially expressed in iNKT cells due to alteration of PLZF function, 17 were found on the PLZF-HDAC7 inverse diagonals and only 4 on the positive diagonals (Fig. 5B-C, labeled genes, Fig. S4C, red asterisks). An additional 4 genes were negatively correlated with HDAC7 function but not differentially expressed during iNKT development (see Table S1), while one was positively correlated. Additionally, Mao, et al. identified BACH2 as a crucial interaction partner of PLZF, and our own data show BACH2 as not differentially expressed but nonetheless as one of the strongest putative upstream regulators of the HDAC7-regulated gene set (Fig. S4A), suggesting that HDAC7 may modulate its targets via a ternary interaction with PLZF. This remarkable degree of overlap strongly supports the idea that HDAC7 is a negative regulator of iNKT cell development that functions at least in part by negatively regulating PLZF-dependent transcription.

Ontologic analysis of HDAC7-regulated genes using Ingenuity Pathway Analysis (IPA) provided strong evidence for their association with both innate-like effector function and inflammatory disease. Canonical pathways associated with the HDAC7-regulated gene set included multiple pathways associated with innate immune signaling and T cell effector function (Fig. S4B, green-shaded pathways, see Table S2 for a complete list of pathways and associated genes), as well as with inflammation and inflammatory disease states (Fig. S4A, blue-shaded pathways), particularly hepatic inflammation. This connection was brought into even sharper relief by two recent GWAS studies of primary sclerosing cholangitis (PSC) and inflammatory bowel disease (IBD), which both identified HDAC7 among the disease-associated loci, and also individually its immediate upstream kinases PKD and SIK2, as well as two isoforms of PKC that are upstream of PKD (Fig. 6A) (Liu et al. 2013, Jostins et al. 2012). Moreover, a remarkably high proportion of the other hits from these studies are downstream of HDAC7, i.e. their expression in iNKT cells is altered by HDAC7ΔP. Of the 176 GWAS risk loci mapping to genes that were expressed in our RNA-seq data, 81 (46%) were regulated by HDAC7 in NKT cells, a much higher degree of overlap than would be expected by chance (P = 3.49 × 10^−16^, binomial distribution) (Fig. 6A). Of the 16 strongest risk loci identified by the Liu, et al. study of PSC, 10 were differentially expressed due to expression of HDAC7ΔP, and 4 more comprised HDAC7 itself, as well as its upstream regulators PRKD2 and SIK2, and also PLZF interaction partner BACH2 (Parra et al. 2005, Mao et al. 2016, Liu et al. 2013) (Fig. 6).

**Figure 6.**
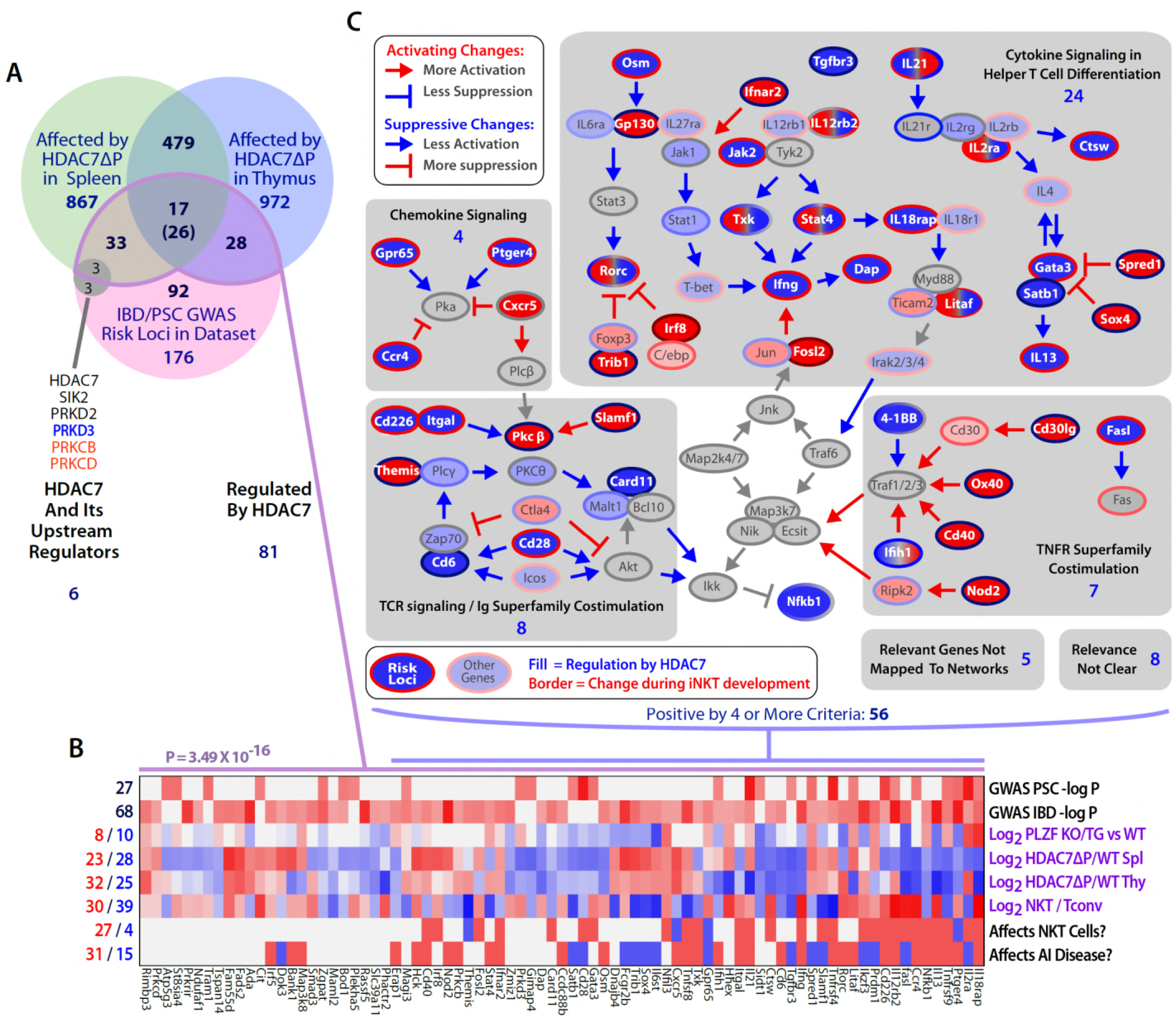
The intersection of HDAC7-regulated genes in iNKT development and GWAS hits for IBD and PSC highlights key signaling pathways. (A) Venn diagram showing enumeration of genes that are GWAS risk loci for PSC and IBD from (Liu et al. 2013, Jostins et al. 2012), and/or also regulated by HDAC7 during NKT development according to our RNA-seq data (FDE > 1.66, P<0.05). The indicated P-value is based on the binomial distribution, using the 13,519 genes scored as expressed under any condition as a basis. (B) Heatmap showing the P-values for the cited GWAS studies in the overlapping set of genes (first two rows), regulation of these genes by PLZF (third row, according to (Mao et al. 2016)), by HDAC7 (rows 4-5), and during normal iNKT development, (row 6, according to Immgen stage-specific data (http://www.immgen.org)), as well as scoring for positive (red) or negative (blue) roles in NK/NKT development/function or autoimmunity, according to literature search (rows 7-8, see Table S1 for citations). (C) Genes that were most relevant to the criteria listed in (B), i.e. positive/significant in four or more measures, were mapped the signaling pathways in which they participate. Shaded areas indicate four distinct, highly populated signaling modules, with number of genes indicated.

To gain a better understanding of the significance of this overlap, we evaluated the 81 risk loci that were regulated by HDAC7 with respect to regulation by PLZF, differential expression in iNKT cells vs. Tconv, functional role in iNKT cell development, and functional role in autoimmune disease (Fig. 6B). This analysis revealed that a large proportion of these genes were functionally important in iNKT development (Fig. 6B, 6^th^ row), while relatively fewer were identified as PLZF targets (Fig. 6B, 3^rd^ row), suggesting that HDAC7 affects autoimmunity and iNKT development via both PLZF-dependent and independent mechanisms. We then further filtered the genes for significance in at least 4 of the 8 criteria examined (Fig 6B), and then manually mapped the resulting 56 genes to their associated signaling pathways (Fig. 6C). Remarkably, all but 13 of these genes could be mapped to one of five interconnected signaling networks, comprising Th1 and Th2 cytokine signaling, chemokine signaling, TCR signaling with its associated costimulatory pathways, and signaling through cell membrane-associated TNF superfamily members (Fig. 6C, dark-colored symbols with white label, gray-shaded areas).

These signaling networks are also heavily populated with HDAC7 targets that were not identified in the GWAS studies (Fig. 6C, light-colored symbols), an observation that is confirmed by IPA analysis of canonical signaling pathways and upstream regulators among HDAC7 targets (Fig. S5A-B). In nearly all cases, HDAC7 regulates these targets in a manner opposite to their regulation during iNKT cell development (Fig. 6C, symbol border vs. fill colors). This regulation by HDAC7 by clearly suppresses downstream signaling in all cases except for TNF superfamily costimulatory signaling, which is potentiated (Fig. 6C, color of arrows per legend). Consistent with our phenotypic findings, nearly half of these genes have positive roles in NK/NKT development/function (Fig. 6B), showing that HDAC7 broadly suppresses several key signaling pathways that are highly important in both NKT cells and in human autoimmune diseases that are similar to the pathology observed in HDAC7ΔP transgenic mice. This remarkable concordance strongly supports the idea that the role of HDAC7 in these cells plays a role in the pathogenesis of PSC and IBD, and identifies a few key signaling pathways as candidates for further interrogation.

### HDAC7 Physically Binds to PLZF and modulates its transcriptional activity

HDAC7 is a class IIA histone deacetylase that lacks intrinsic DNA binding capacity and requires binding to target transcription factors to modulate transcription at specific loci (Yang and Seto 2008). Class IIA HDACs typically act as dominant corepressors, as in the case of MEF2, which is converted from a transcriptional activator to a repressor upon class IIA HDAC binding (McKinsey et al. 2000). PLZF belongs to the BTB-ZF family of transcription factors (Beaulieu and Sant'Angelo 2011) previously reported to interact with class IIA HDACs (Verdin, Dequiedt, and Kasler 2003, Chauchereau et al. 2004); indeed, one group has even demonstrated in vitro and in vivo binding of HDAC7 to PLZF in a separate cell type (Lemercier et al. 2002). This suggested that HDAC7 might modulate PLZF activity in thymocytes through direct physical binding. Determining if this is the case directly is somewhat challenging, however, as the abundance of PLZF in wild-type thymocytes is very low, being restricted to a small population of iNKT precursors. To circumvent this difficulty, we made cell lysates from PLZF-transgenic thymocytes and immunoprecipitated them with antibodies to endogenous HDAC7. These experiments showed a specific interaction between HDAC7 and PLZF in thymocytes (Fig. 7A).

**Figure 7.**
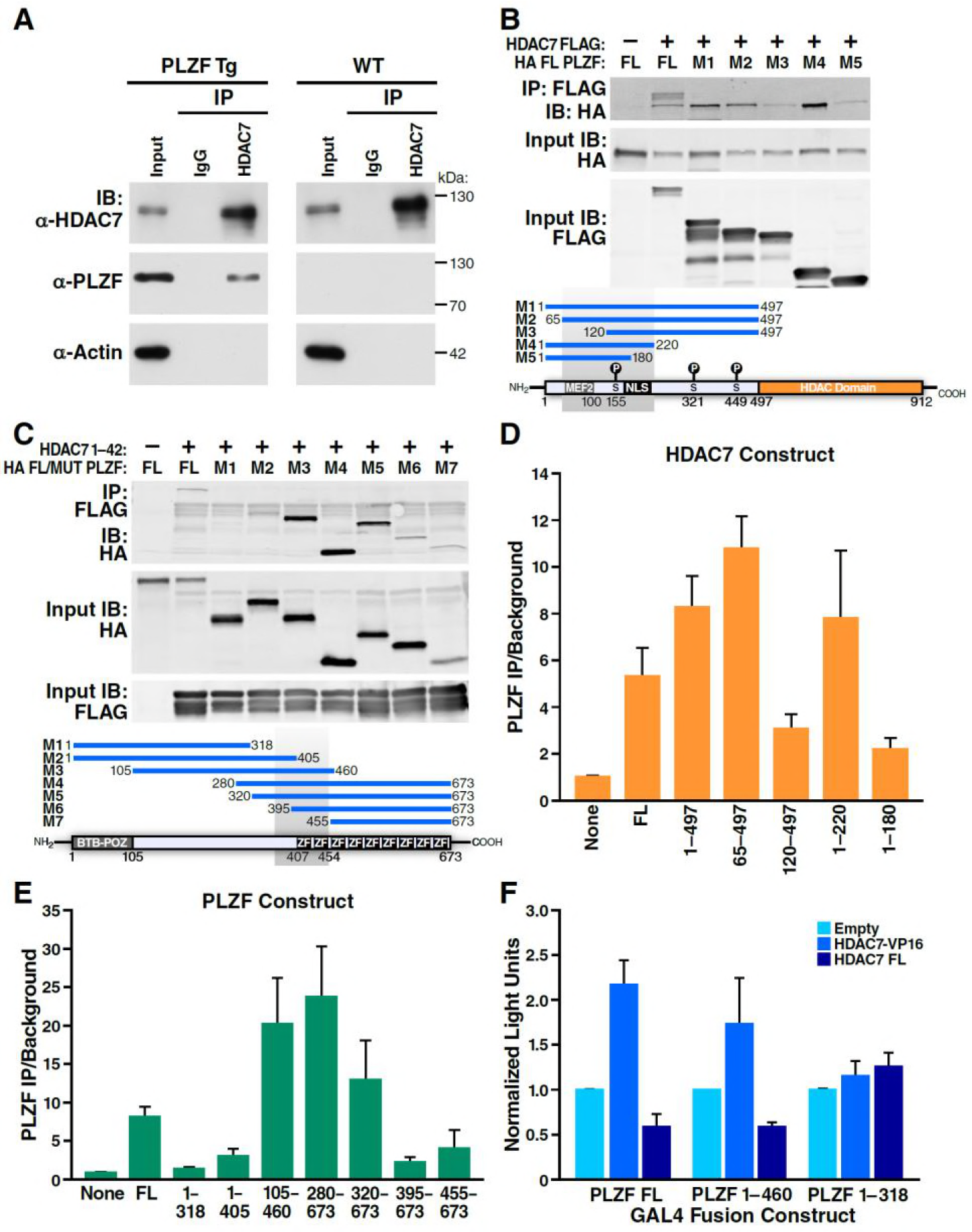
HDAC7 Can Physically Bind and Functionally Antagonize PLZF Transcriptional Activity. (A) Immuno-blots showing co-immunoprecipitation with endogenous HDAC7 of PLZF from PLZF-transgenic thymocytes. (B) Immunoblot showing Co-immunoprecipitation of HA-tagged full-length PLZF from transfected 293T cells with the indicated FLAG-tagged truncation mutants of HDAC7 (C) Immunoblot showing Co-immunoprecipitation of HA-tagged PLZF truncations as indicated, with the FLAG-tagged HDAC7 1-497 (D), (E) Quantification of Immunoprecipitated protein/input protein for the pairs of constructs in (B) and (C) respectively. Ratios shown are normalized to the background signals for each individual experiment. Error bars indicate SEM of 4-7 individual experiments for each pair of constructs. Shaded areas in diagrams in (B) and (C) indicate areas defined as required for interaction based on this analysis. (F) Firefly luciferase activity from 293T cells transfected with a Gal4(5)/SV40 minimal promoter reporter construct, normalized to *Renilla* luciferase values from an EFlα promoter-driven reporter construct. In addition to the reporters, cells were transfected with constructs encoding the Gal4 DNA-binding domain (1-142) fused to the indicated segments of PLZF, as well as empty vector, full-length HDAC7, or HDAC7 1-497 fused to the HSV VP16 transcriptional activation domain (410-490). Error bars represent SEM of four individual experiments.

To further define this interaction, we co-transfected FLAG-tagged full-length or truncated HDAC7 with full length HA-tagged PLZF (Fig. 7B, D), or conversely different truncations of PLZF with the (interacting) HDAC7 N-terminal adapter domain (residues 1-497, Fig. 7C, E). After Immunoprecipitation of transfected lysates with anti-FLAG agarose beads, we quantified the amount of PLZF protein pulled down vs. input levels over 3-6 separate experiments for each construct, using the LiCor Odyssey system (Fig. 7D-E). The results of this analysis identify residues 65-200 of HDAC7, containing the MEF2-interacting domain through the first PKD phosphorylation site, as the interacting region (Fig. 7D). Analysis of the PLZF deletions identified a region from residues 320-450, encompassing a proline-rich tract and the first two zinc finger domains, as critical for interaction (Fig. 7E).

Although the precise mode of transcriptional regulation by PLZF remains unclear, with different domains exhibiting activating and repressive activity in varying contexts (Sadler et al. 2015, Puszyk et al. 2013, Melnick et al. 2002), we next wanted to examine if HDAC7 physical binding to PLZF could modulate its transcriptional activity. We transfected 293T cells with fusions of the GAL4 DNA binding domain (residues 1-142) to full-length PLZF, an HDAC7-interacting mutant of PLZF (1-460), or a non-interacting mutant (1-318), together with a SV40 minimal promoter-Gal4(5)-firefly luciferase reporter and an EF-1α promoter-driven Renilla luciferase reporter (Fig. 7F, see Fig. S5 for a diagram). To these constructs were added empty vector (Fig. 6F, light blue bars), a vector encoding full-length HDAC7 (Fig. 7F, dark blue bars), or one encoding a fusion of the HDAC7 1-497 interacting domain with the VP16 transcriptional activation domain (HDAC7-VP16, Fig. 7F, medium blue bars). Measurement of luciferase activity in lysates from these cells showed that co-transfection of FL HDAC7 with FL PLZF or the 1-460 truncation reduced transcription from the Gal4-luc construct, while it did not affect transcription when co-transfected with the non-interacting 1-318 mutant (Fig. 7F). Conversely, HDAC7-VP16 increased transcription from the interacting PLZF constructs but not the non-interacting one (Fig. 7F). These experiments, together with our characterization of the HDAC7-PLZF interaction and transcriptional targets above, provide strong evidence that in thymocytes HDAC7 regulates PLZF in the same manner as MEF2 and other transcription factors, functioning as a TCR signal-dependent co-repressor that helps to silence PLZF-associated promoters in the absence of appropriate signals. This mechanism is highly likely to account for at least part of the effect on HDAC7 on innate effector differentiation.

### Restoring iNKT Cells Ameliorates Tissue-Specific Autoimmunity

We earlier reported that HDAC7ΔP mice develop spontaneous tissue-specific autoimmunity, with about 80% developing obliterative exocrine pancreatitis and concomitant T-cell infiltration in stomach, liver and small intestine within eight months (Kasler et al. 2012). Although this had been previously attributed solely to a defect in negative selection of conventional thymocytes, the striking absence of iNKT cells in HDAC7ΔP mice spurred us to consider whether disrupted innate effector development might also contribute to this autoimmune syndrome. Indeed, the very tissues vulnerable to T-cell infiltration in HDAC7ΔP mice, notably the small intestine, liver and hepatobiliary mucosa, are typically populated by PLZF-dependent innate effectors such as iNKT and mucosal-associated invariant T (MAIT) cells (Fan and Rudensky 2016). We thus set out to determine if restoring iNKT cells could alter the course of HDAC7ΔP–induced autoimmunity.

In our earlier studies, we found that that HDAC7ΔP-mediated autoimmunity is dominantly transferable in mixed BM chimeras if a 5-fold excess of HDAC7ΔP-derived bone marrow is used. While engraftment at these ratios produced comparable populations of WT and HDAC7ΔP Tconv in peripheral tissues, we did not assess the reconstitution of the iNKT compartment in those studies (Kasler et al. 2012), leaving open the possibility that there was an uncharacterized recessive component to the autoimmunity. Attempts to adoptively transfer mature iNKT cells directly into HDAC7ΔP mice failed to effectively restore tissue-resident iNKT populations (Fig. S6A-C). Instead, we generated two sets of hematopoietic chimeras to determine if restoring iNKT cells using Vα14 bone marrow could ameliorate disease compared to WT bone marrow (Fig. 8A). When irradiated recipients were reconstituted with a 1:5 mixture of Vα14: HDAC7ΔP bone marrow, peripheral iNKT cells were effectively rescued to normal levels, while in recipients receiving a 1:5 WT: HDAC7ΔP mixture they were still essentially absent (Fig. 8B).

**Figure 8.**
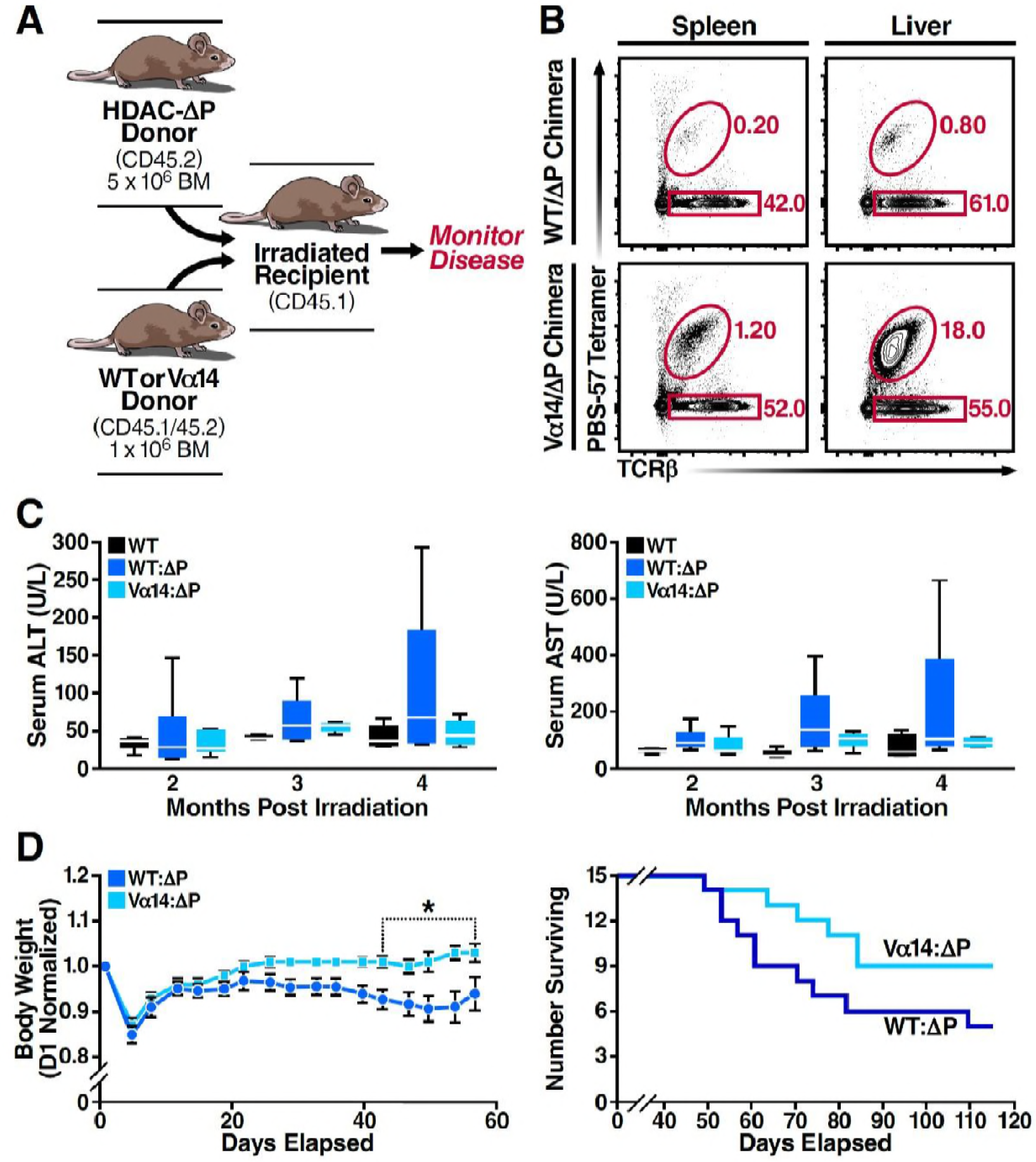
Loss of PLZF-Dependent Innate Effectors in HDAC7ΔP Contributes to Tissue-Specific Autoimmunity. (A) Schematic of mixed BM chimeras used to monitor *HDAC7ΔP*-mediated autoimmunity time course and severity. Lethally irradiated CD45.1 Boy/J recipients were reconstituted (6×10^6^ cells) with a 1:5 mixture of either WT(CD45.1): *HDAC7ΔP*(CD45.2) or Vα14(CD45.1): *HDAC7ΔP*(CD45.2) bone marrow cells. (B) *Vα14* bone marrow (bottom) robustly restores peripheral iNKT cells (Tet^+^ TCRβ^+^ in liver and spleen in mixed BM chimeras, while WT bone marrow does not. Plots are representative of two sets of independently made chimeras. (C) Plasma concentration of liver (ALT, AST) markers of tissue damage over time measured in WT mice compared to *Vα14*: *HDAC7ΔP* and WT: *HDAC7ΔP* BM chimeras. (D) Body weight (left) and survival (right) of mixed BM chimeras over time post-irradiation. Weights in (D) were normalized to starting weight on Day 1 post-irradiation and measured twice a week thereafter. Survival (D, right) was assessed by monitoring for spontaneous death twice a week or by euthanasia after reaching a clinical endpoint of at least 20% body weight loss compared to peak weight post-irradiation. Using Kaplan-Meier analysis, p=0.0616 by Gehan-Breslow-Wilcoxon tests. Bars on graphs indicate mean ± SEM (error bars); whiskers on box-and-whiskers plots represent min to max. Data in (C) were collected from N=6 mice per group; data in (D) and (E) were combined from three independent experiments with N=16 mice total per group. Statistical significance in (D) was determined using two-way ANOVA; *p ≤ 0.05. Bonferroni post-tests were used for pairwise comparisons.

Comparing these cohorts over time, we noted Vα14: HDAC7ΔP chimeras had significantly lower peak plasma levels of ALT and AST, commonly used as an indication of liver damage, than WT: HDAC7ΔP chimeras (Fig. 8C). Both cohorts eventually perished from exocrine pancreatitis and had similar pancreatic lipase levels in plasma (Fig. S6D), yet Vα14: HDAC7ΔP chimeras exhibited significantly improved body weight maintenance in the first two months post-engraftment (Fig. 8D, left) and a reduced overall mortality rate (Fig. 8D, right) compared to WT: HDAC7ΔP chimeras. These results provide evidence that disruptions in innate effector development, particularly the loss of iNKT cells in the hepatobiliary tract, exacerbates tissue specific autoimmunity in the HDAC7ΔP setting. Restoring this missing innate effector population resulted in enhanced survival and a significant reduction in the severity of disease.

## Discussion

### HDAC7 Nuclear Export is an essential Licensing Step in Innate Effector Development

The discovery and characterization of innate effector lymphocytes has transformed our understanding of T-cell receptor signaling, barrier protection at mucosal surfaces, and the evolutionary origins of the vertebrate immune system, yet the identification of key regulatory factors that control naïve versus innate effector development in thymocytes has remained elusive. We demonstrate here that the epigenetic regulator HDAC7 serves as a gatekeeper of this developmental fate decision in the thymus. When HDAC7 is prevented from releasing its genomic targets in response to TCR stimulation, PLZF-dependent innate effector development is blocked and iNKT cells become diverted to a naïve-like fate, characterized by lack of expression of memory or NK markers and a failure to produce effector cytokines. Conversely when HDAC7 function is lost, naïve development is reduced and more thymocytes develop as CD8-expressing innate effectors. Thus, appropriately regulated nuclear export of HDAC7 is an important licensing step that permits both negative selection and the acquisition of alternative cell fates, such as PLZF-dependent agonist selection to the iNKT lineage.

In this study, we focused on iNKT cells due to their relatively high abundance and easy identification using CD1D tetramers, but we suspect that HDAC7ΔP similarly abrogates development of other PLZF-dependent innate effector subtypes, including rare MR1-restricted MAIT cells and γδ NKT cells (Chandra and Kronenberg 2015, Fan and Rudensky 2016). In contrast, another well-described innate effector type, CD8αα+ IELs localized in small intestine (Mayans et al. 2014), are only slightly reduced in HDAC7ΔP mice (data not shown), consistent with their PLZF-independent derivation (Cheroutre, Lambolez, and Mucida 2011). The identification of a committed precursor to innate lymphoid cells that transiently expresses high amounts of PLZF (Constantinides et al. 2014) also raises the intriguing possibility that development of these cell types may be regulated by class IIA HDACs as well. Furthermore, the main mechanism of action we investigate here, HDAC7 antagonism of PLZF via direct interaction, may be generalizable to other members of the BTB-POZ-ZF family. For example, the signature transcription factor of Tfh cells, Bcl6, is known to associate with HDAC4 (Lemercier et al. 2002) (Crotty 2014). A class IIA HDAC/BTB-ZF axis may thus regulate T cell or ILC development at additional branch points. Additionally, in recent years a number of transcriptional regulators and epigenetic modifiers – including Jarid2, NKAP, HDAC3, and Ezh2 (Pereira et al. 2014, Thapa et al. 2013, Dobenecker et al. 2015) – have been identified that regulate iNKT ontogeny. At least one member, HDAC3, physically associates with class IIA HDACs as part of a larger co-repressive complex (Fischle et al. 2002). Devising systems to investigate these relationships as well as HDAC7 association with PLZF via ChiP-Seq and other genomic-scale approaches is a current priority in our laboratory.

### HDAC7 Control of iNKT Cell Development Modulates the Susceptibility of Liver to Autoimmune Attack

By restoring the missing iNKT population with the use of Vα14 donor bone marrow, we significantly attenuated the severity and time course of HDAC7ΔP-mediated autoimmune liver disease, resulting in improved liver function, better body weight maintenance, and reduced overall mortality. Although specific rescue of iNKT cells did not provide protection in all tissues – almost all Vα14: HDAC7ΔP chimeras eventually developed the same ultimately lethal exocrine pancreatitis as WT: HDAC7ΔP chimeras – our studies nonetheless reveal an important new role for impaired iNKT development as an exacerbating factor in liver autoimmunity. Since both HDAC7 and PLZF influence the development of several non-iNKT innate effector subtypes that would not have been restored with Vα14 bone marrow, it is tempting to speculate that restoring these other subsets might ameliorate tissue destruction and T-cell infiltration due to HDAC7ΔP in other organs.

Innate effector T-cells are often considered frontline first-responders to infection that amplify and orchestrate the early immune response to invading pathogens. Thus, it was somewhat surprising to uncover a protective or anti-inflammatory role for iNKT cells in attenuating tissue destruction. Additional studies will be required to uncover the mechanisms through which iNKT cells provide protection, but for now we favor a model in which innate effectors occupy tissue niches at their sites of residence, limiting access of other immune cells into those sites. In HDAC7ΔP mice, escape of autoreactive Tconv due to impaired negative selection may produce a potentially but not necessarily pathogenic population, which requires the additional loss of PLZF-dependent innate effectors from their target tissues to create an opening for infiltration. This “two-hit” model may explain multiple types of tissue-specific autoimmunity, in which genetic lesions that generate excess self-reactive lymphocytes are coupled with separate or related defects in tissue-resident innate effector populations at specific sites, rendering these tissues particularly vulnerable to attack.

Our findings likely hold considerable relevance to understanding the etiology and mechanisms contributing to some types of human autoimmunity. Indeed, common variant single nucleotide polymorphisms (SNPs) in the HDAC7 gene are significantly associated with human autoimmune and auto-inflammatory diseases, namely primary sclerosing cholangitis (Liu et al. 2013) and inflammatory bowel disease (Jostins et al. 2012). Additional common variant SNPs in kinases known to export Class IIA HDACs via phosphorylation, including SIK2 and PRKD2, are also associated with primary sclerosing cholangitis (Liu et al. 2013), suggesting aberrant regulation of HDAC7 nuclear export as a causative mechanism. Moreover, the genes that we identified as regulated by HDAC7 in iNKT development show a striking overlap with other risk loci from these GWAS studies (Fig. 6A), suggesting that the broad HDAC7 regulatory network may be a crucial nexus that underlies susceptibility to several autoimmune diseases of considerable clinical importance. Indeed, mapping the overlapping GWAS loci to their associated signaling networks revealed a remarkable clustering around a few important signaling pathways in iNKT and effector development, including IL12, IL21, IL18, IFNG, and IL4, as well as Ig- and TNF-superfamily costimulatory pathways. Deciphering the complex relationship between HDAC7, PLZF and other HDAC7 interaction partners, the observed modulation of these pathways, and the resulting cellular and pathologic phenotypes will be a major task for us going forward. We hope that this effort will illuminate the way forward in translating our finding that reestablishing missing iNKT cells can ameliorate HDAC7-mediated hepatic autoimmunity into potential therapeutic modalities for the analogous human diseases, based on the restoration of innate effector function.

## Materials and Methods

### Study Design

The initial objective of this work was to investigate the molecular mechanisms behind the control of iNKT development by HDAC7, which was an observation we made incidentally in our prior characterization of the general role of HDAC7 in thymic T cell development. The idea that HDAC7 might do this at least in part via interaction with PLZF arose from the review of older literature on these molecules showing they interact. The notion that the role of HDAC7 in iNKT cells has a bearing on the tissue distribution of autoimmunity due to altered HDAC7 function arose from the concordance between NKT-populated tissues and those showing disease in HDAC7ΔP transgenic mice. This idea was highlighted in importance by the publication of GWAS studies after the initiation of our work that statistically associated HDAC7 and its regulatory network with human diseases affecting the same tissues. We investigated these questions using a combination of cell culture and transgenic mouse models in which the function of HDAC7 and/or PLZF was altered in thymocytes. Parameters measured include cellular abundance in different tissues, T cell effector function after *ex-vivo* stimulation, luciferase expression, protein-protein interactions, global transcript abundance, and various clinical measures associated with autoimmune disease, as detailed in the following sections.

With the exception of our RNA-seq study, which was done in one experiment using three biological replicates for each condition, all results depicted in this work are based on at least two completely independent trials, comprising at least 3 biological replicates. Larger sample sizes than this were used as feasible, based on the availability of experimental genotypes of interest, prospective estimates of the statistical power required to show significance for robust effects, and the constraints of time and resources required for analysis. No data that were collected were excluded from the study unless there was clear evidence of a technical failure in data collection, or in the case of the animal studies, morbidity/mortality that was clearly unrelated to the pathologic conditions under study. Except where otherwise indicated in the figure legends, all control-experimental pairs were composed of sex-matched littermates, and all primary immune phenotypes were measured in animals between 5 and 8 weeks of age.

### Mouse Strains and procedures

All mice were housed in a specific pathogen-free barrier facility at the Gladstone Institutes, in compliance with NIH guidelines and a UCSF IACUC animal use protocol. All experimental strains were on a C57BL/6 (B6) genetic background. B6, BoyJ, Vα14/Jα18 transgenic (Tg(Cd4-TcraDN32D3)1Aben) and PLZF transgenic (C57BL/6-Tg(Cd4-Zbtb16)1797Aben/J) mice were obtained from Jackson Laboratories. Mice deleting HDAC7 or expressing the *HDAC7ΔP* transgene under the control of the p56^lck^ proximal promoter were prepared as described elsewhere (Kasler et al, 2011) (Kasler et al. 2012). Hematopoietic chimeras were prepared as follows: Recipients (8-10 wk-old BoyJ or BoyJ X B6) mice were irradiated with a split dose of 700 + 500 Rads, 3 hrs. apart, from a ^137^Cs source (J.S. Shepherd & Associates). Mice were reconstituted with 5X10^6^ bone marrow cells from WT (Boyj or B6 X BoyJ heterozygote), HDAC7ΔP Tg (CD45.2), HDAC7 KO (CD45.2), or Vα14/Jα18 (CD45.2) transgenic donors, injected retro-orbitally in 200 μl of PBS. Bone marrow cell suspensions were prepared by crushing tibias and femurs, dissociating marrow cells in PBS, and purifying mononuclear cells by Ficoll gradient centrifugation. Serum for AST/ALT/lipase analysis was collected by tail vein incision and analyzed by the UCSF Clinical Laboratory at SFGH.

### Flow cytometric analysis of immune cell populations

Cell suspensions were prepared from mouse thymus and spleen by crushing, dissociation of cells by pipetting, straining through 40 micron nylon mesh, and ACK lysis. Cells were prepared from liver using an additional Percoll gradient centrifugation. For analysis of cytokine expression, cells were cultured for 4 hours post-isolation with 50ng/ml PMA (Sigma) plus 0.5μM ionomycin (Sigma) and for 1 hour with 0.5μg/ml Brefeldin A (Sigma) prior to staining. Viability staining was performed for 15 minutes in the dark at room temperature using eFluor 506 or eFluor 780 fixable viability dyes (eBioscience) at 1:1000 in PBS. Surface staining with CD1D tetramers and fluorochrome-conjugated antibodies was performed for 30 minutes on ice in PBS with 2% FCS, followed by either fixation in PBS/1% PFA or fixation/permeabilization with the eBioscience FOXP3 intracellular staining kit. Intracellular staining for cytokines or transcription factors was performed for 1 hour on ice in eBioscience FOXP3 Perm/wash buffer. Analytical flow cytometry was performed using a BD FACS Calibur DxP or LSRII. Data processing for presentation was done using FlowJo 10.0 (Treestar Inc.). Cell sorting was performed using a BD FACS-Aria. CD1D-αGalCer tetramers (PBS-57), conjugated with either phycoerythrin (PE) or allophycocyanin (APC) were obtained from the NIH tetramer core (http://tetramer.yerkes.emory.edu/). The following commercial antibodies were used for flow cytometry: CD11a-PE-Cy7 (eBioscience), clone M17/4; CD18-PE (eBioscience), clone M18/2; CD24-PE-Cy7 (BD Bioscience), clone M1/69; CD3-APC-EF780 (eBioscience), clone 2C11; CD4-BV650 (BD Bioscience), clone RM4-5; CD4-PE (BD Bioscience), clone GK1.5; CD4-APC (BD Bioscience), clone RM4-5; CD44-PE-Cy7 (eBioscience), clone IM7; CD44-APC-Cy7 (BD Bioscience), clone IM7; CD44-APC (eBioscience), clone IM7; CD45.1-Pacific Blue (eBioscience), clone A20; CD45.1-FITC (eBioscience), clone A20; CD45.2-V500 (BD Bioscience), clone 104; CD45.2-PE-Cy7 (BioLegend), clone 104; CD5-APC (BD Bioscience), clone 53-7.3; CD62L-APC-Cy7 (BD Bioscience), clone MEL-14; CD69-PE (eBioscience), clone H1 2F3; CD8-Alexa 700 (Tonbo), clone 53-6.7; CD8-PerCP (biolegend), clone 53-6.7; CXCR3-PE (eBioscience), clone cxcr3-173; Eomes-PE (eBioscience), clone Dan11mag; Ly6C-APC (eBioscience), clone hk1.4; NK1.1-PE-Cy7 (BD Bioscience), clone pk136; NK1.1-APC-Cy7 (BD Bioscience), clone pk136; PLZF-PE (eBioscience), clone Mags.21f7; T-bet-PE-Cy7 (eBioscience), clone ebio4b10; TCRβ-PerCP-5.5 (BD Bioscience), clone H57-597; TCRβ-APC-Cy7 (BD Bioscience), clone h57-597; TCRγδ-APC (BD Bioscience), clone GL3; Vg6.3/6.2-PE (BD Bioscience), clone 8f4h7b7.

### RNA-seq analysis of gene expression

Cell suspensions were prepared from thymus and spleen of 6-8 week old wild type B6, Vα14/Jα18 transgenic, or Vα14/Jα18 X *HDAC7ΔP* mice. iNKT cells were sorted by FACS using antibodies to TCRβ(+) and the PBS-57 CD1D-αGalCer tetramer(+). Naïve Tconv were sorted using antibodies to CD4(+), CD8(−), TCRβ(+), and CD44(−). Cells (250,000-2,000,000) were purified from three littermate triplets for each strain (18 samples total), and total RNA (200ng to 4μg) was prepared using the Rneasy Plus Mini Kit (Qiagen). Double-stranded cDNA libraries were prepared by the Gladstone Institutes Genomics Core using the Nugen Ovation kit. The Libraries were sequenced by the UCSF Center for Advanced Technology using the Solexa HiSeq 4000 instrument. Six barcoded samples were loaded per lane. FASTQ files (approximately 5.5X10^7^ reads each) were mapped to the UCSC Mouse genome Build 37 (Mm.9) using Bowtie2 (Johns Hopkins University). Approximately 4X10^7^ (~75%) of reads per sample were mapped uniquely to the mouse genome. Gene-level tabulation, quality control, and expression analysis was done on •SAM format files generated by BOWTIE2 using SeqMonk 0.33 (http://www.bioinformatics.babraham.ac.uk/projects). Ontologic analysis and pathway mapping were performed using Ingenuity Pathway Analysis (http://www.ingenuity.com/). All primary data associated with these experiments have been deposited at GEO (https://www.ncbi.nlm.nih.gov/geo, accession GSE105026), and a summary of all gene expression data and statistics for differentially expressed genes is provided in Table S1.

### Plasmids, transfections, and reporter assays

The human PLZF coding sequence (RCAS(B)-Flag-PLZF), deposited by Peter Vogt (Shi and Vogt 2009) was obtained from Addgene. N-terminally HA-tagged full-length PLZF was amplified from this coding sequence using the following Primers: N-terminal PLZF Bam HI, HA tag, EcoRV, Bsa BI, Hpa I: 5’ aaaaaaggatccacc atg tat ccc tac gat gtt cca gat tat gcg ata tca atc gtt aac atg gat ctg aca aaa atg gg; C-terminal Swa 1, stop, Not 1: 5’ cct cta cct gtg cta tgt gtg att taa atgattagataagcggccgcaaaaaa 3’. This amplification product was subcloned into pCDNA3.1(+) using BamH1 and Not1 sites. Different PLZF truncations were amplified from this construct and sub-cloned into the introduced flanking sites (further details on request). For the Gal4 DNA-binding domain-PLZF fusion constructs, the Gal4 DNA-binding sequence was amplified from a plasmid template and inserted into the Eco RV sites of full-length or truncated PLZF expression constructs described above. Construction of full-length human HDAC7 and HDAC7-VP16 fusion-encoding expression vectors is described elsewhere (Dequiedt et al. 2003). Other truncated, FLAG-tagged HDAC7 constructs were amplified from these templates and religated into pCDNA3.1(+). The Gal4(5) SV40-Firefly luciferase reporter construct was prepared by ligation of an oligonucleotide cassette containing 5 GAL4 recognition sites into the Sma 1 site of pGL2 Promoter (Promega).

For pulldown experiments, 10cm dishes seeded the previous day with 3.2X10^6^ HEK 293T cells were transfected with 20μg of total DNA, consisting of 10μg each of PLZF and HDAC7 constructs or the corresponding empty vectors, using CaPO_4_/chloroquine. After 48 hours, cells were harvested for interaction analysis. For reporter assays, 6-well dishes seeded with 0.8X10^6^ HEK 293T cells/well were transfected using CaPO_4_/chloroquine with 6.1μg of total DNA, consisting of 2 μg each of gal4(5) luc, gal4-PLZF fusion construct, and empty vector or HDAC7 expression construct, plus 100ng of EF1α *Renilla* luciferase. Cells were harvested for luciferase assay 48 hours after transfection, and luciferase activity was measured using the Promega Dual-Luciferase assay kit.

### Co-immunoprecipitations and western blots

For the co-immunoprecipitation of endogenous HDAC7 with transgenic PLZF in thymocytes, thymocyte lysates from wild-type and PLZF transgenic mice were prepared using p300 lysis buffer (250 mM NaCl, 0.1% NP-40, 20 mM NaH2PO4, pH 7.5, 5 mM EDTA, 30 mM sodium pyrophosphate, 10 mM NaF, and HALT protease/phosphatase inhibitors (Thermo Fisher Scientific). After clarification (5 minutes, 13,000Xg) and pre-clearing (3 hours at 4°C with proteinA/G agarose beads), lysates were immunoprecipitated with either 1μg/ml of α-HDAC7 antibody (H-273, Santa Cruz Biotechnology) or 1μg/ml of rabbit IgG isotype control antibody (Cell Signaling) at 4°C overnight. The lysates were then incubated with 50μl of protein A/G agarose beads (Santa Cruz Biotechnology) at 4°C for 4 hours, followed by washing 5 times with p300 lysis buffer. Immunoprecipitated proteins from the beads were eluted with non-reducing Laemmli SDS PAGE sample buffer by boiling for 3 min. For pulldown analysis of HDAC7-PLZF truncation mutants, 10cm dished were harvested and lysed in 0.8mL of P300 buffer, clarified by spinning 5 min. at 13,000g, then incubated for 4 hours at 4°C with 30μL/sample of FLAG M2-agarose beads (Sigma). After 4 washes with p300 buffer, bound proteins were eluted from the beads by addition of 100μL of reducing Laemmli SDS-PAGE sample buffer, followed by a 5-minute incubation at 95°C.

After SDS PAGE and transfer to nitrocellulose, membranes were probed with antibodies against HDAC7 (H-273, Santa Cruz Biotechnology), PLZF (D9, Santa Cruz Biotechnology), and β-actin (Abcam), HA epitope (Cell Signaling), or FLAG epitope (Sigma), overnight at 4°C. After washing and incubation with HRP- or IRDye- (LiCor) conjugated antibodies, signal was detected using chemiluminescence and film or a LiCor Odyssey scanner respectively. Bands for quantitative pulldown analysis were quantified from the scanner output using ImageJ (Wayne Rasband, National Insititutes of Health).

## Acknowledgments

We thank G. Maki and T. Roberts for figure preparation, M. Cavrois, M. Maiti, A. Uebersohn for technical assistance, C. Doherty for coordinating plasma lipase and liver panel measurements, A. Abbas, A. Chawla, D. Sheppard, and members of the Verdin laboratory for helpful comments and discussion, and M. Ott and J. Roose for critically reading the manuscript.

---

**Figure S1.**
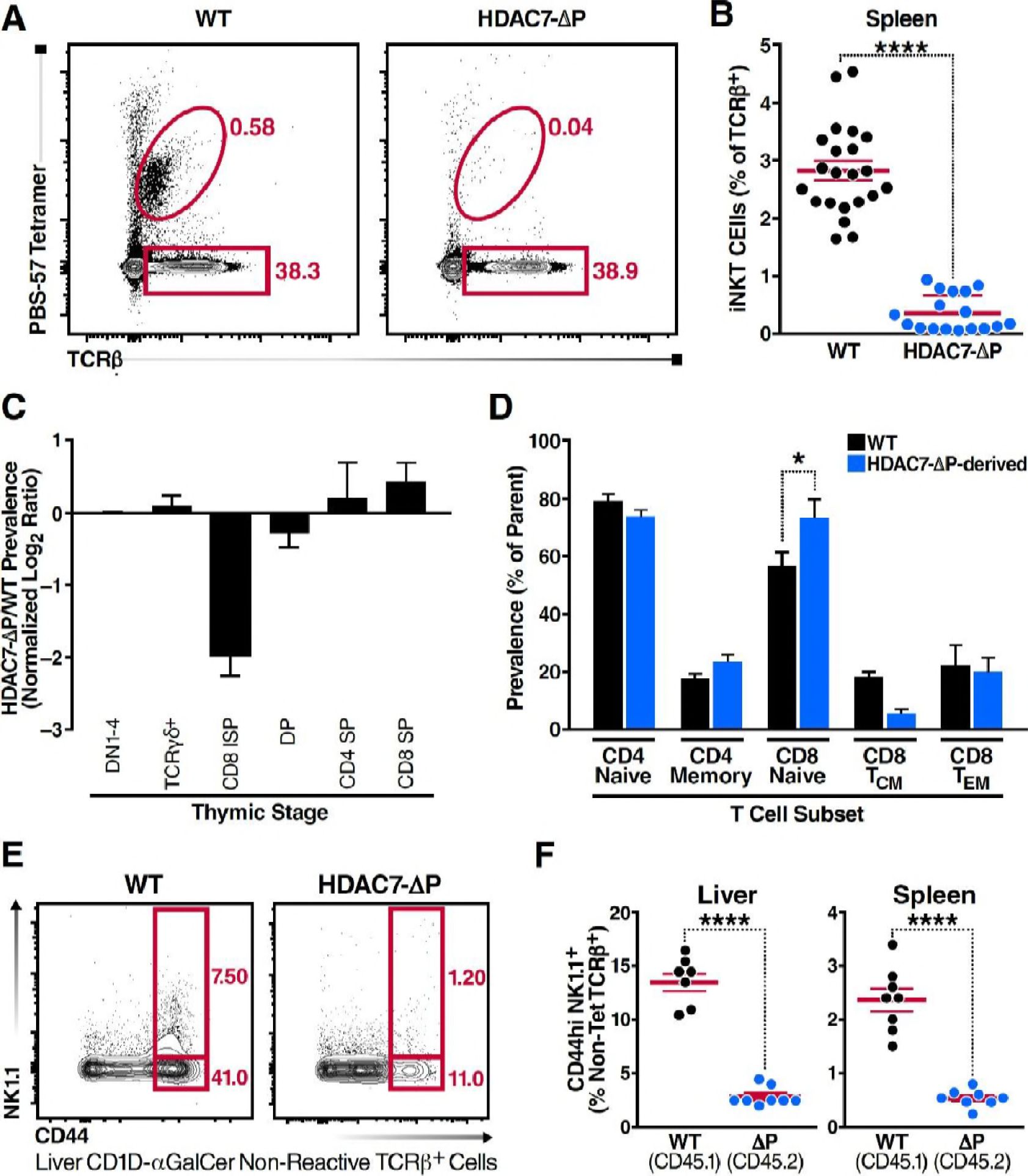
HDAC7-ΔP Arrests Thymic iNKT Development, Related to Figure 1. (A, B) Representative flow cytometric plots (A) and total quantification (B) of iNKT cells and conventional αβ T-cells from spleen. Data in (B) are combined from 8 independent experiments. (C) Log_2_ fold ratio of HDAC7ΔP-derived (CD45.2) to WT-derived (CD45.1) cells at the indicated thymocyte stages in mixed bone marrow chimeras. A composite DN1-4 engraftment ratio (Lin^−^CD4^−^CD8^−^) was calculated per mouse to normalize the ratio at each successive stage. (D) Proportion of T-cell subsets plotted as percentage of parent from HDAC7ΔP-derived (CD45.2) or WT-derived (CD45.1) bone marrow in irradiated chimeras. CD4 naïve are defined as CD44^lo^CD4^+^, CD4 memory as CD44^hi^CD4^+^, CD8 naïve as CD44^lo^CD62L^hi^CD8^+^, CD8 central memory (T_CM_) as CD44^hi^CD62L^hi^CD8^+^, and CD8 effector memory (T_EM_) as CD44^hi^CD62L^lo^CD8^+^. (E, F) Representative flow cytometric plots (E) from liver and total quantification (F) from liver and spleen of type II iNKT cells (Tet^−^ CD44^hi^ NK1.1^+^ TCRβ^+^) in mixed bone-marrow chimeras depending on bone marrow of origin. Data in (B) are combined from a subset of experiments described in Fig 1D-E, with N=8 per group. Statistical significance in (F) was determined using a 2-tailed Student’s T test. P<0.0001 for both WT-HDAC7ΔP comparisons.

**Figure S2.**
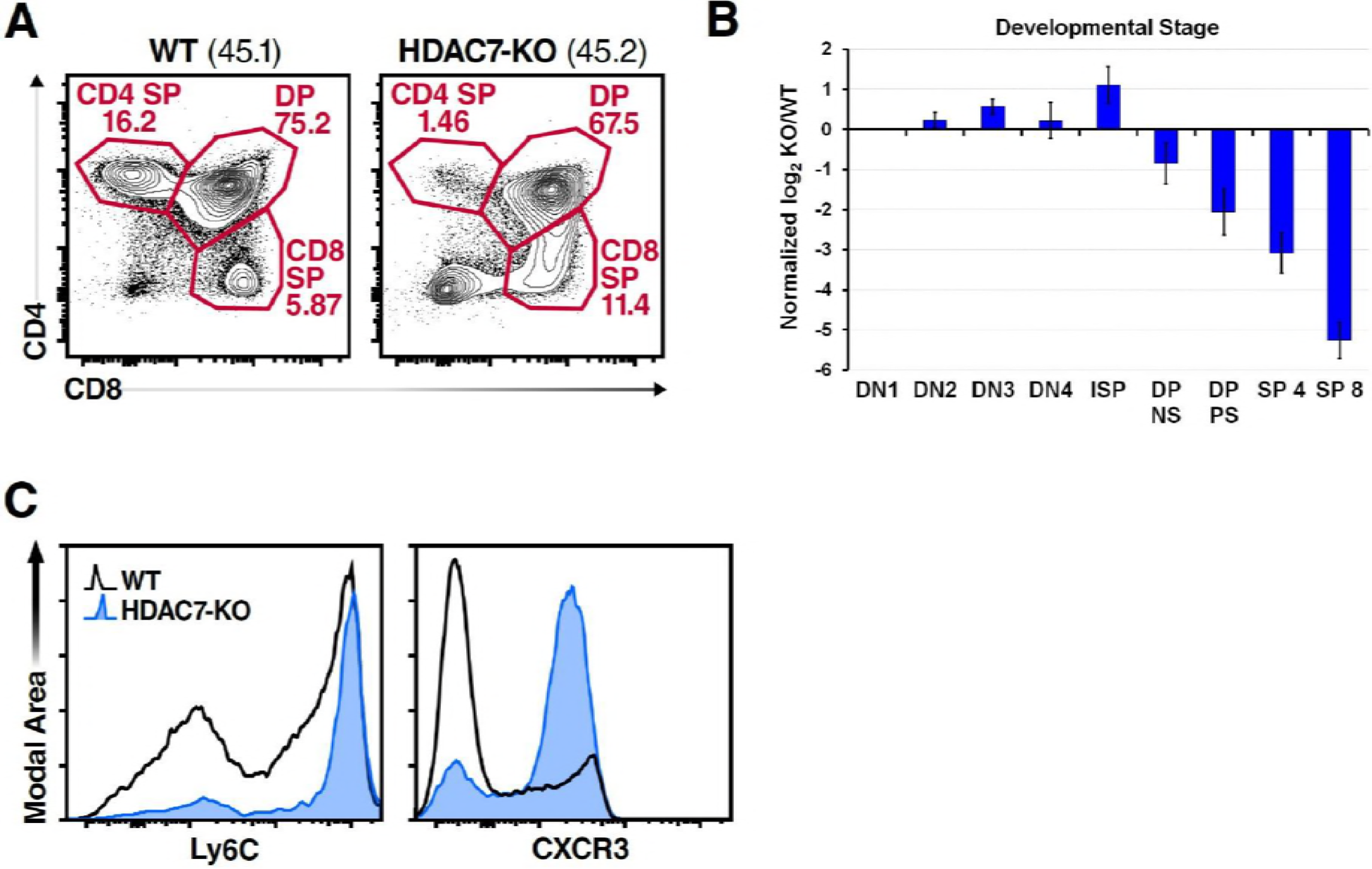
Expansion of Innate Memory CD8 T-Cells in HDAC7-KO Mice, Related to Figure 2. (A) Representative flow cytometric plot of total thymocytes from WT-derived (CD45.1) and HDAC7 KO-derived (CD45.2) bone marrow in irradiated mixed bone-marrow chimeras. (B) Quantification of Log_2_ ratios of KO/WT Cells at each of the indicated thymic developmental stages. Results are shown for 4 individual chimeric mice, +/− std. dev. (C) Surface expression of Ly6C and CXCR3 from peripheral CD8 T-cells. Black unfilled histograms correspond to WT, blue-filled to *HDAC7 KO*. Plots represent 3 independent experiments with 2-4 mice per group.

**Figure S3.**
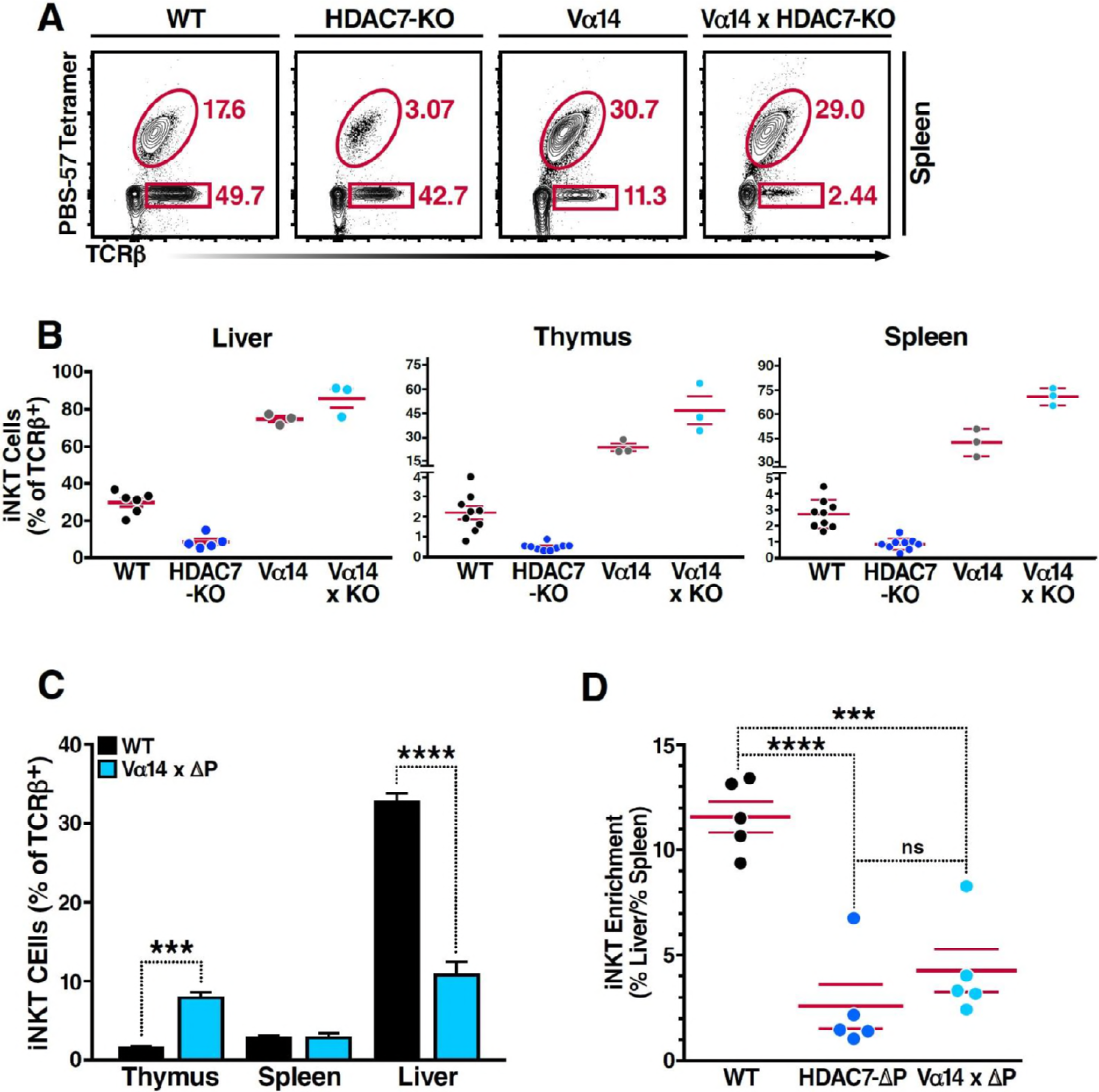
HDAC7-AP iNKT cells Fail to Accumulate in Liver, Related to Figure 3. (A, B) Restoration of iNKT cells (Tet^+^ TCRα^+^) in *HDAC7 KO* mice by expression of the Vα14-Jα18 TCRα transgene. Representative plots for the indicated genotypes shown in (A) with total quantification shown in (B) for liver (left), thymus(middle), and spleen (right) for 3 pairs of littermate mice. (C) Proportion of iNKT cells expressed as percent of total TCRβ^+^ T-cells in thymus, spleen and liver from WT and Vα14 × HDAC7ΔP mice. (D) Fold enrichment of iNKT cells in liver (% total TCRβ^+^) over spleen (% total TCRβ^+^) in WT and Vα14 × HDAC7-ΔP mice. Statistical significance was determined using unpaired two-tailed t tests; ***p ≤ 0.001, ****p ≤ 0.0001.

**Figure S4.**
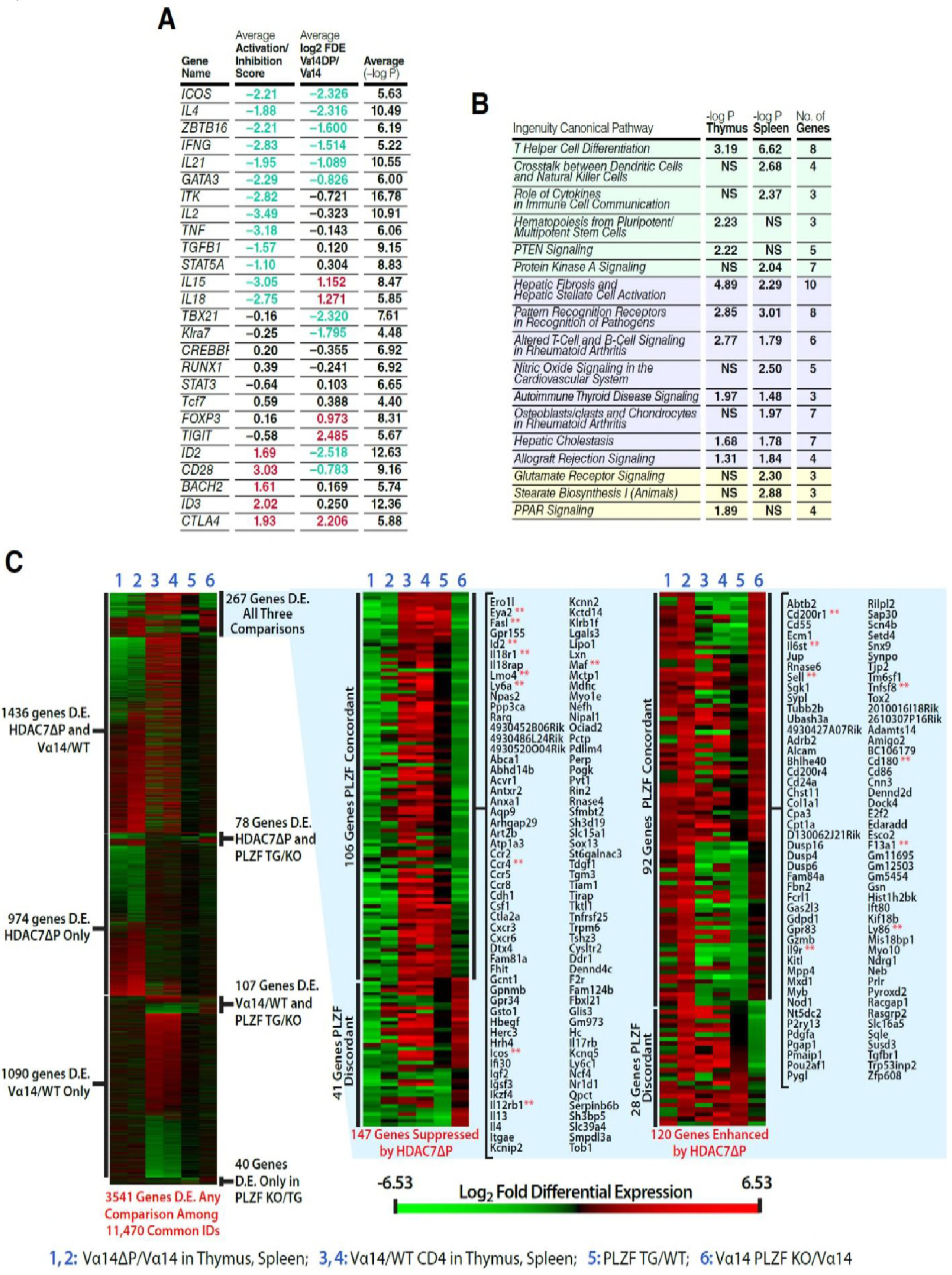
HDAC7 Regulates a Cassette of Genes in Glycolipid-Reactive Cells That is Highly Relevant to Innate Effector Function, Inflammation, Autoimmunity, and Autoimmune Liver Disease. Related to Figure 5. (A) Table showing top putative upstream regulators of the sets of genes from Fig. 5A that were suppressed by *HDAC7ΔP* in thymus or spleen. Column 2 shows the average activation or inhibition z-score of the indicated genes, based on IPA analysis of differential regulation of their targets in the data. Column 3 shows the average log2 fold differential expression of the indicated genes in *Vα14* X *HDAC7ΔP* vs *Vα14* Tg thymocytes and splenocytes. (B) Table showing IPA top canonical signaling pathways overrepresented in the sets of genes from (A) that were suppressed by HDAC7ΔP in thymus or spleen. Red-shaded pathways refer to inflammation or inflammatory disease states, blue-shaded pathways to pathways involved in innate effector differentiation or function. (C) Heat maps showing differential expression values (red is upregulated, green downregulated) for the 11,470 genes sharing unique common IDs between our RNA-seq data and data on the role of PLZF in iNKT cell development recently published in (Mao et al. 2016). Columns 1-2 show the effect of HDAC7ΔP expression in PBS-57 tetramer-reactive *Vα14* × *HDAC7ΔP* vs *Vα14* Tg thymocytes (1) and splenocytes (2) from our RNA-seq data. Columns 3-4 show the differential expression of the same genes in *Vα14* vs. naïve CD4SP Tconv in thymus (3) and spleen (4), also from our data. Columns 5-6 show differential expression of these genes in PLZF transgenic vs. wild-type CD4SP thymocytes (4), or in *Vα14* X *Luxoid* (PLZF null) vs. *Vα14* (5) PBS-57 tetramer-reactive CD4SP thymocytes, taken from published data (Cohen et al. 2013, Mao et al. 2016), archived at GEO (accession GSE81772). The block at left shows differential expression values for 3,541 genes among the 11,470 common IDs that are >1.67-fold differentially expressed in at least one of the comparisons shown. The heat maps at center and right show 267 genes that are differentially expressed due to HDAC7ΔP expression (1, 2) and during NKT development (3, 4), and also due to altered PLZF function (5, 6). Identities are shown for 198 genes that are suppressed by HDAC7ΔP and enhanced by PLZF (center block), or conversely enhanced by HDAC7ΔP and suppressed by PLZF (right block). Genes found to be bound by PLZF (Mao et al. 2016) are indicated by double red asterisks.

**Figure S5.**
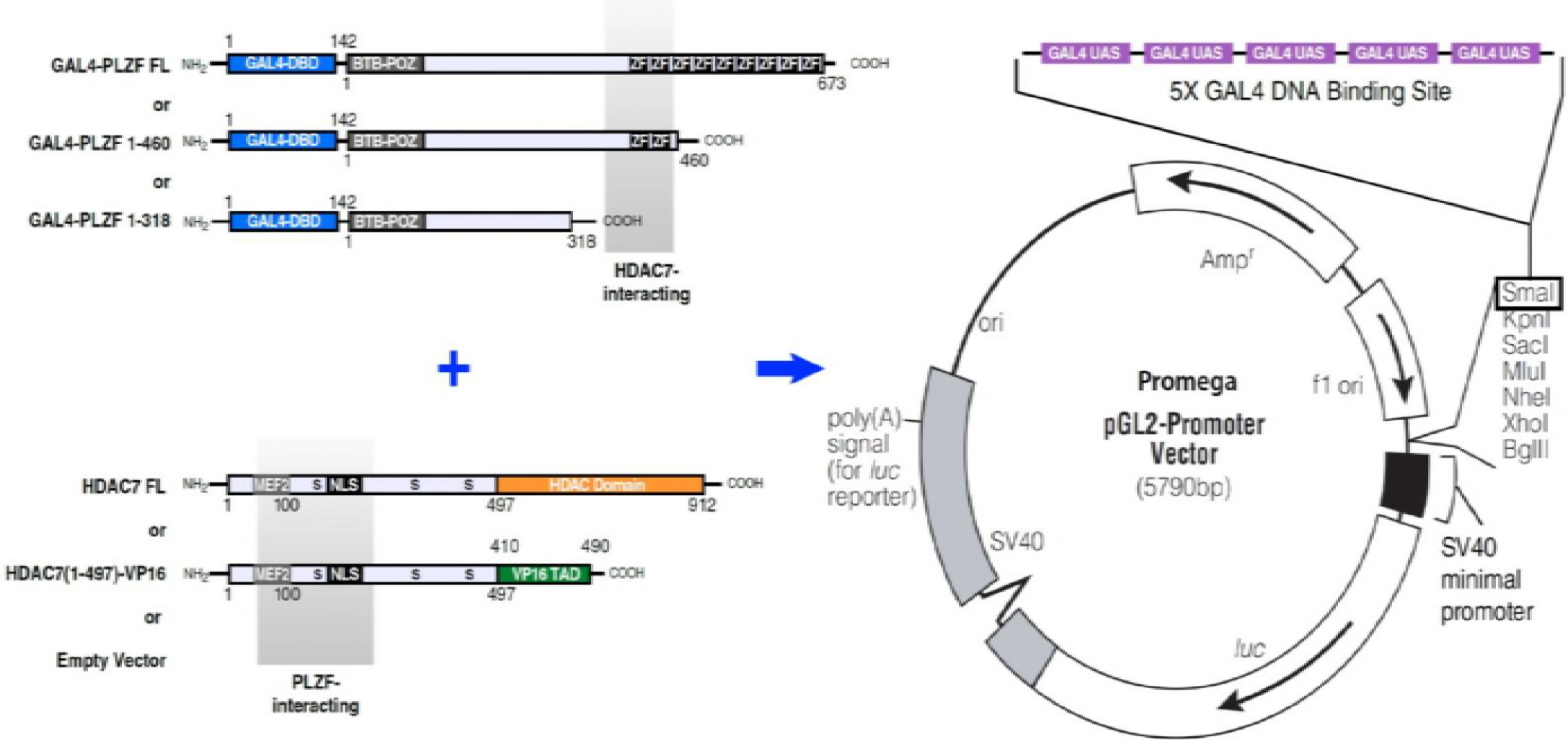
Diagram of GAL4-PLZF, HDAC7, and GAL4 reporter constructs employed in experiments shown Figure 7F in main text. The GAL4 DBD-PLZF fusion and HDAC7 constructs shown were co-transfected into HEK293T cells, together with the 5XGAL4 site-containing pGL2 Promoter luciferase reporter shown at right.

**Figure S6.**
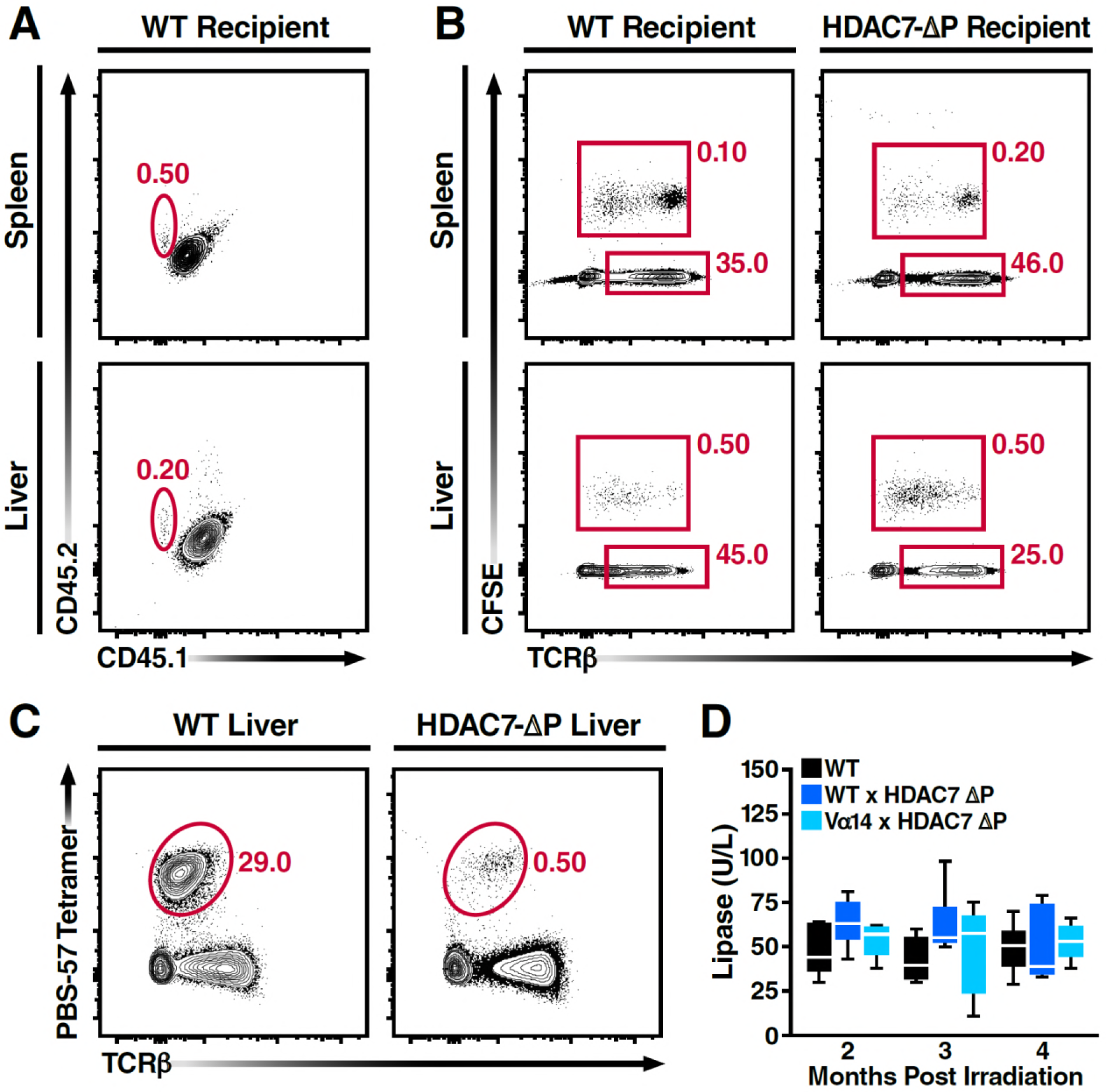
Restoration of iNKTs with Vα14-Jα18 bone marrow alters autoimmune disease course, related to Figure 8. (A) Representative flow scatters showing CD45.1 vs. CD45.2 expression in spleens and livers of WT CD45.1/.2 heterozygote recipients, three days after retro-orbital transfer of CD45.2 5×10^6^ iNKT cells. (B) Identification of CFSE-labeled adoptively transferred T-cells in liver and spleen of WT and HDAC7-ΔP in mice from (A). (C) Representative flow scatters showing, TCRβ^+^, PBS-57 tetramer-reactive cells in Livers of mice from (A). Transferred iNKT cells were isolated from spleens and livers of Vα14-Jα18 transgenic mice and enriched to 85+% before transfer. Data are representative of two independent experiments, N=3 mice per group total. (D) Plasma concentration of lipase over time in WT mice versus Vα14-Jα18 / ΔP and WT / ΔP BM chimeras. Samples were obtained from a subset of the cohorts described in Fig 7D-E, with N=6 mice per group.

**Notes Accompanying Supplementary Table 1.** Normalized RNA expression values shown were generated from SAM format mapped read files using SeqMonk v0.33 (www.bioinformatics.babraham.ac.uk/projects/), employing their RNA-seq quantification pipeline. Relative expression values for expressed genes were further median-centered before analysis of differential expression. Genes included in the table were filtered based on these expression values (at least one sample > 2.0).

